# Lamin B2 follows lamin A/C- mediated nuclear mechanics and cancer cell invasion efficacy

**DOI:** 10.1101/2020.04.07.028969

**Authors:** Marina Vortmeyer-Krause, Mariska te Lindert, Joost te Riet, Veronika te Boekhorst, Rene Marke, Ramanil Perera, Philipp Isermann, Tom van Oorschot, Monika Zwerger, Fengwei Yang, Martin Svoreň, Anotida Madzvamuse, Jan Lammerding, Peter Friedl, Katarina Wolf

## Abstract

Interstitial tumor cell invasion depends upon complex mechanochemical adaptation of both cell body and the rigid nucleus in response to extracellular tissue topologies. Nuclear mechanics during cell migration through confined environments is controlled by A-type lamins, however, the contribution of B-type lamins to the deformability of the nucleus remains unclear. Using systematic expression regulation of different lamin isoforms, we applied multi-parameter wet-lab and in silico analysis to test their impact on nuclear mechanics, shape regulation, and cancer cell migration. Modulation of lamin A/C and B2 but not B1 isoforms controlled nuclear deformation and viscoelasticity, whereby lamin B2 generally followed lamin A/C-mediated effects. Cell migration rates were altered by 5 to 9-fold in dense collagen environments and synthetic devices, with accelerated rates after lamin downregulation and reverse effects after lamin upregulation, with migration rates strongly depending on nuclear shape change. These findings implicate cooperation of lamin B2 with lamin A/C in regulating nuclear mechanics for shape adaptation and migration efficacy.

**Summary:** Nuclear deformability during cancer cell invasion and metastasis is critically regulated by lamin A. Here, researchers showed that lamin B2 also contributes to nuclear mechanics, implicating cooperating lamin networks regulating nuclear integrity, migration efficacy, and metastatic tumor progression.

## Introduction

Cell migration along or through complex three-dimensional (3D) tissues is an essential process during tissue formation, homeostasis and remodeling, however, also underlies tumor invasion and metastasis. Moving cells are confronted with connective tissue of varying topology and density, including loose fibrillar networks adjacent to densely packed, highly rigid stroma (Butcher et al., 2009; Charras and Sahai, 2014; Provenzano et al., 2008; Weigelin, 2012). Cells respond to these extracellular matrix (ECM) topologies and geometric confinements by either matrix metalloproteinase (MMP)-dependent proteolytic ECM degradation, or by reversibly deforming their shape (Wolf et al., 2003). To accommodate such shape changes, the ellipsoid nucleus, which is 2-10 times stiffer compared with the surrounding cytoplasm (Lammerding, 2011), undergoes complex deformation and transiently adapts elongated, hourglass-like or other, irregular shapes during cell passage through dense ECM. When the capacity of the nucleus to deform further is exhausted, it represents the rate-limiting organelle during cell migration (Davidson et al., 2014; Fu et al., 2012; Harada et al., 2014; Wolf et al., 2013).

The deformation kinetics of the cell nucleus in 3D environments is determined by its viscoelastic behavior, which can be modeled as a combination of springs and dashpots, and which collectively determine its deformability under mechanical forces. Elasticity (*syn*. stiffness, rigidity) describes how much the nucleus instantaneously deforms under force and reversibly bounces back into its original shape after force release, not unlike a rubber band.Viscosity describes the time-dependent behavior in response to force application, and often involves creep, i.e. continuous deformation under a constant load. The viscoelastic mechanical behavior of the nucleus depends on contributions from both the chromatin and the nuclear lamina, with chromatin primarily governing behavior for small deformations and the lamina dominating the mechanical resistance at larger extensions (Pajerowski et al., 2007; Stephens et al., 2017).

The nuclear lamina consists of a layer of interconnected intermediate filaments below the double nuclear membrane, forming a mechanosensitive cortical network, as well as nucleoplasmic filaments inside the nucleus (Kolb et al., 2011). The cortical filaments are formed by proteins of the lamin family, including A-type (A, C) and B-type (B1, B2) lamins which assemble into homotypic filaments and antiparallel polymers (Ho and Lammerding, 2012). While B-type lamins are expressed in all soft proliferating cells, A-type lamins become upregulated upon cell maturation and differentiation (Broers J.L.V., 2014; Constantinescu et al., 2006). Accordingly, A-type lamins were identified as primary mediators of nuclear stability and elasticity, using a range of experimental strategies including microaspiration, mechanical strain, microfluidic setups or atomic force spectroscopy (AFS) on adherent or suspended cells as well as on isolated nuclei (Davidson et al., 2014; Harada et al., 2014; Kaufmann et al., 2011; Lammerding et al., 2006; Lammerding et al., 2004; Lange et al., 2017; Lange et al., 2015; Lautscham et al., 2015; Schape et al., 2009; Swift et al., 2013). In addition, lamin A expression, often measured as lamin A:B ratio, positively correlated with ECM rigidity (Swift et al., 2013), indicating lamin A as the main structure mediating resistance to deformation force. Accordingly, migrating rates of nontransformed fibroblasts and invasive cancer cells in confining substrates in vitro, such as collagen, PDMS channels or transwell chambers, depend on the expression level of lamin A/C (Davidson et al., 2014; Greiner et al., 2014; Harada et al., 2014; Infante et al., 2018; Lautscham et al., 2015; Ribeiro et al., 2014; Rowat et al., 2013; Shin et al., 2013; Yadav et al., 2018), in line with findings that softer tumor cells associate with higher invasion rates (Swaminathan et al., 2011). With cancer progression, the de-differentiation of tumor cells and acquisition of stem cell properties is often accompanied by de-regulated A- as well as B-type lamin expression (Alhudiri et al., 2019; Broers J.L.V., 2014; Chow et al., 2012; Denais and Lammerding, 2014; Saarinen et al., 2015; Wazir et al., 2013). However, whereas the relevance of lamin A/C in regulating nuclear deformability and cell migration has been established, the role of B-type lamins is less clear. By combining biomechanical, functional and in silico analyses, we here address how both A- and B-type lamins control nuclear mechanics, and how their up- and downregulation translates to concordantly altered migration efficacy of invasive cancer cells. We identify lamin B2, in addition to the known functions of A-type lamins, as critical regulator of the viscoelasticity of the nucleus as well as of nuclear shape change and effective migration in intact cells. These data provide an inventory for complementary experimental techniques and models to dissect differential roles of lamin isoforms in nuclear mechanobiology and cell migration in environments of different dimensionality and complexity.

## Results

### Lamin expression regulation and distribution

Invasive HT1080 fibrosarcoma cells were used for probing lamin functions on nuclear mechanics and the capacity to migrate in 3D collagen (Krause et al., 2013; Wolf et al., 2013). HT1080 cells expressed lamin isoforms A/C, B1 and B2, which distributed around the chromatin in a rim-like fashion, and formed tube-like invaginations, here visible as intranuclear linear signal and speckles (Figs. 1A and B). HT1080 cell variants with down- or upregulated lamin A, C, B1 or B2 were generated and compensatory expression regulation by the other, non-targeted lamin isoforms was generally not detected (Figs. 1B and C; S1A). Transient downregulation for 48 hours by a pool of small inhibitory (si) RNAs (knockdown; RNAi) resulted in 11-40% remaining protein levels in a lamin-dependent manner (Figs. 1B and D; S1B), and constitutive overexpression increased lamin protein levels by 2.3- to 4.5-fold (Figs. 1C and D). These changes of lamin isoform expression were confirmed on the single-cell level by immunofluorescence against specific lamin isoform, with weak but homogeneous residual lamin distribution at the nuclear rim after lamin downregulation (Figs. S1A and C). Conversely, lamin upregulation induced higher fluorescent signals at the nuclear lamina, as well as more prominent membrane foldings or intranuclear tubes, with the exception of lamin C, which appeared to concentrate rather towards intranuclear tubes than the rim (Fig. S1D). Together, combining transient dowregulation and permanent upregulation in the same HT1080 cell source resulted in an 8-27 fold change from lowest to highest expression of the respective lamin isoforms (Fig. 1D).

**Figure 1.**
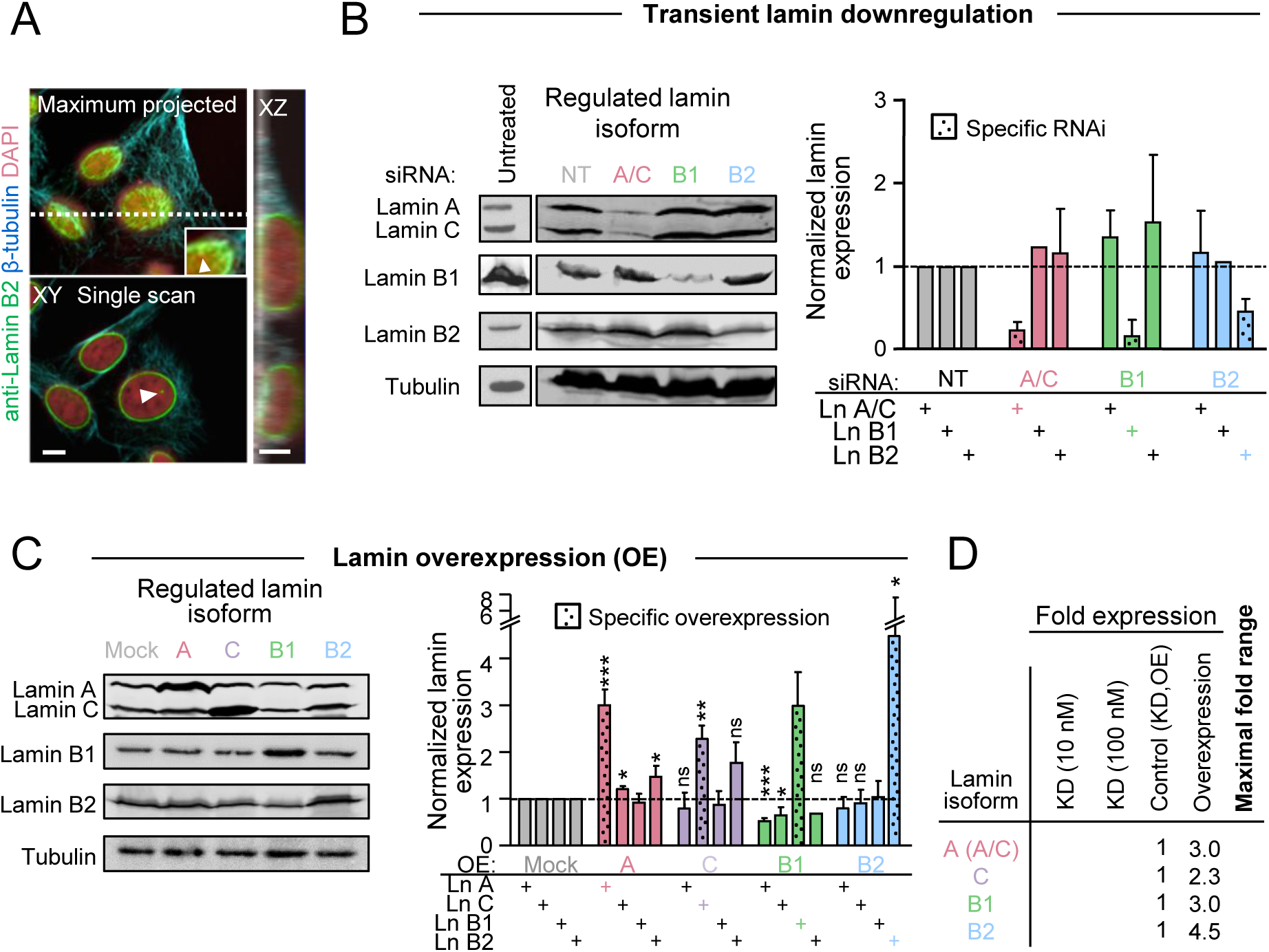
Transient knockdown or stable overexpression of lamin isoforms in HT1080 cells. **(A)** Lamin localization in cells shown in XY and XZ presentation. Upper left panel; maximal Z-projection to show overall lamin content. Lower left; single, central slice to visualize lamin ring from nuclear rim, and linear and dot-like lamin signals indicating tube-like invaginations (arrowheads). Right panel; XZ representation, created along white dotted line from upper left image for vertical view. Colored staining as indicated. Image bars: 5 μm. **(B,C)** Modification of lamin levels after transient knockdown (48h; 10 nM RNAi) or stable overexpression of specific lamin isoforms. From lamin-specific bands from the western blots (left), respective signal intensity was calculated by densitometry. Lamin intensity rates were normalized to both control (non-targeting control, NT; Mock) signal and respective tubulin loading control, and normalized controls are shown as 1 (graph; right). Lamin isoform-specific bands after indicated modification and respective matching antibody were quantified and shown as bars with dots and marked in color below the graph. Bars and whiskers are mean +SD/SEM. **(B)** N = 2-6; **(C)** n = 2-5. For statistical analyses, two-tailed unpaired Students t-test was used; ***, P ≤ 0.001; **, P ≤ 0.01; *, P ≤ 0.05; ns, non significant. **(D)** Mean lamin regulation level (deviating from 1 for NT or Mock) after indicated modulation of lamin isoforms. Data are a summary from Figs. 1, S1B, and not shown (n = x). KD, knockdown.

### Control of cell and nuclear mechanics by different lamin isoforms

To quantify and compare the impact of A- and B-type lamins on the deformability of a cell in response to transient compression, AFS combined with confocal microscopy was applied (Fig. 2A). We positioned the bead-functionalized cantilever on top of the nucleus of live HT1080 cells attached to plastic dishes during cell culture, and transiently compressed (‘approached’) the cell (Fig. 2B; Video1). To calculate the response of cell and nucleus to deformation relevant for cell migration in confinement (see Figs. 3A,B; 5) (Krause et al., 2013), we applied a broad force range between 1 and 1000 nN on adherent cells. These yielded nuclear deformations from 0.5 to ∼4 μm as a linear function, with a plateau reached at 200 nN and 4 μm indentation, from which small intendations were used for the calculation of the cells elastic modulus E (Fig. 2C). After lamin A/C down- or lamin A upregulation, elasticity values changed to 0.27 and 0.77 kPa (as compared 0.41 and 0.47 kPa in control cells; Fig. S2). Cell and nuclear stiffness thus was reduced to 0.66-fold or increased to 1.62-fold, respectively, when compared to normalized (1-fold) control values (Fig. 2D, Table 1). Similarly, the elastic modulus after lamin B2 downregulation was reduced 0.75-fold and after upregulation increased 1.43-fold, whereas no significant change was noted after deregulation of lamin B1 or C. Using 20 nN compression force, indentation after depletion of lamin A/C increased from 2.9 to 3.25 μm (thus by 12%), and to 3.1 after lamin B2 downregulation (thus by 7%), but remained again unchanged after lamin B1 pertubation (Fig. 2E). Similar softening was noted when forces up to 150 nN were used (Fig. 2F). Thus, deep (3-4 µm) and more subtle (0.5 µm) indentation yielded comparable results for a function of lamins A/C and B2 in nuclear deformability. In line with these data, regulation of nuclear deformation by lamin A/C using microfluidics, aspiration, nuclear strain or AFS experimentation previously yielded 1.5 to 4-fold changes (Davidson et al., 2014; Harada et al., 2014; Kaufmann et al., 2011; Lammerding et al., 2006; Lange et al., 2015), whereas the impact of lamin B2 deregulation on nuclear stiffness was not investigated.

**Table 1.**
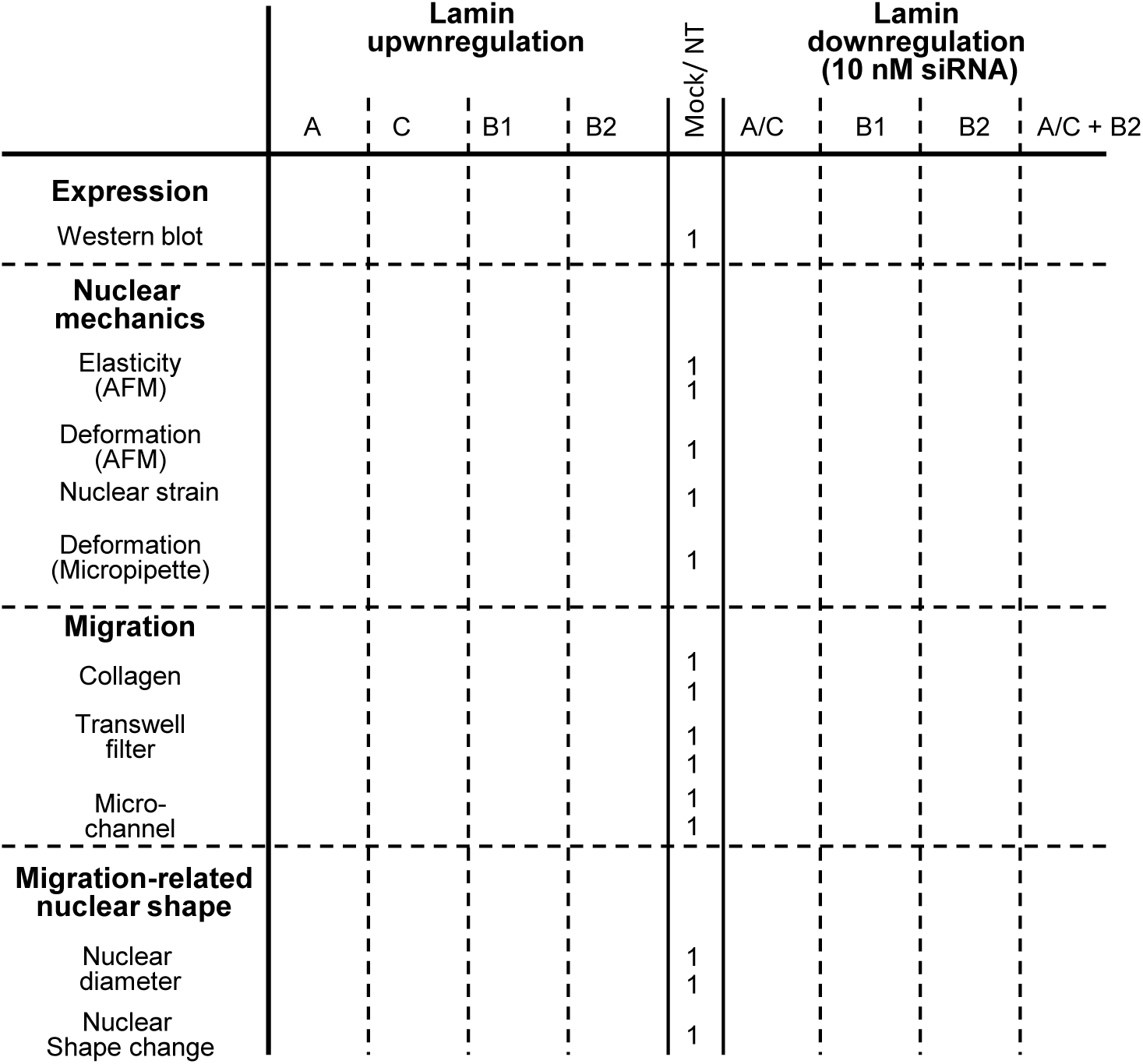
Summary of all measured lamin-related changes shown in Figures 1-7. Displayed numbers are fold changes as compared to NT or Mock control in HT1080, MDA-MB231, MV3 and MCF-7 cells. Note that almost exclusively the fold changes for lamin B2 are closer to 1 than for lamin A/C, meaning that all effects mediated by lamin B2 are present but somewhat weaker than lamin A/C.

**Figure 2.**
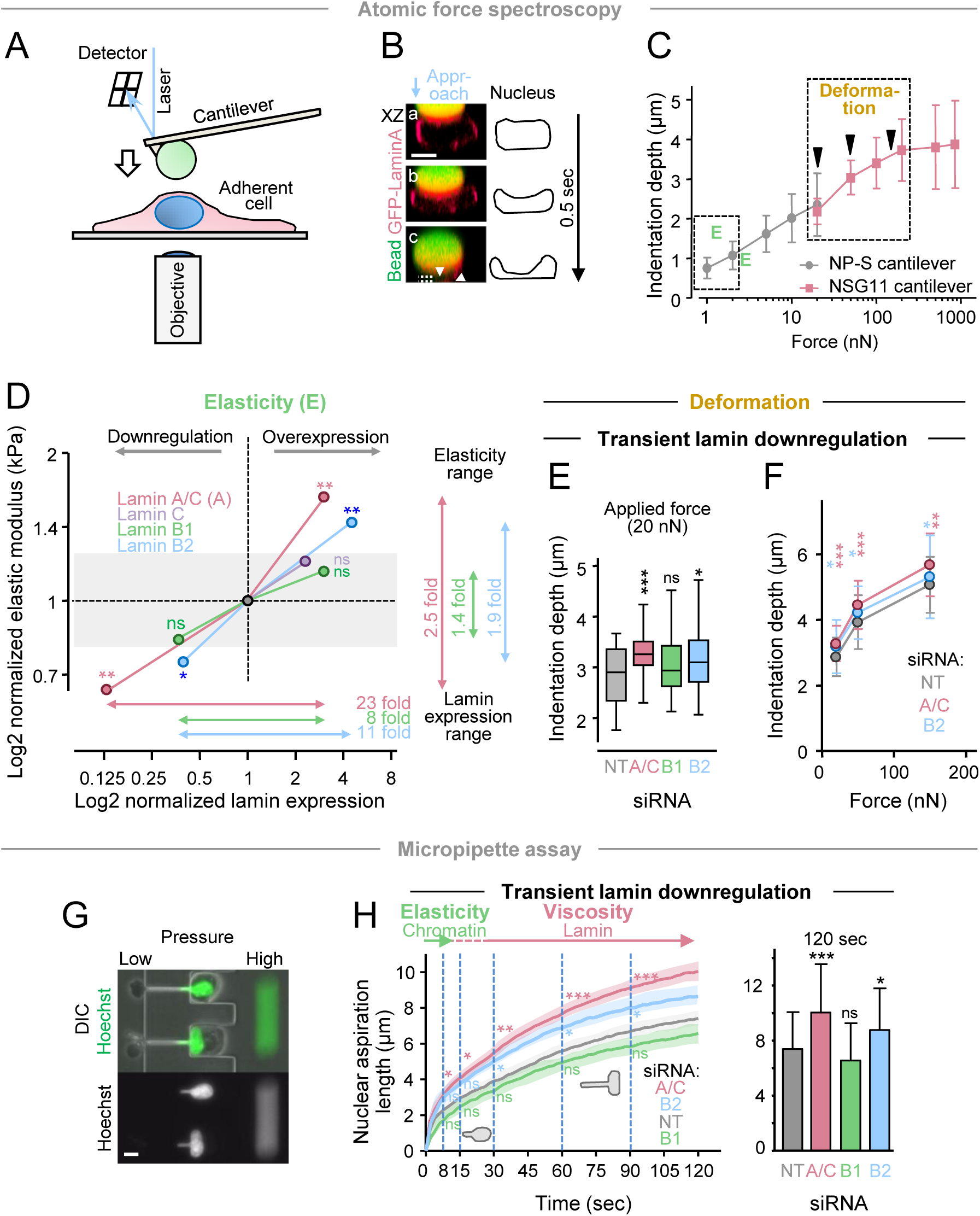
Mechanics of tumor cell nuclei correlates with expression of lamins A/C and B2. **(A)** Depiction of AFS methodology. **(B)** Visualization of nuclear compression of an adherent cell upon probing by a bead-coupled cantilever. XZ image sequence (corresponding to Video 1) of HT1080 cell stably expressing GFP-lamin A before (panel a) or during (panel b-c) application of 700 nN force by a cantilever-coupled bead, with dotted lines and white arrowheads indicating respective nuclear heights. Numbers, time in seconds. Of note, compression was carried out at slower speed and with higher contact force, in order to visualize the measurement principle. Image bar, 10 mm. **(C)** Nuclear penetration as a function of contact force measured by bead modified cantilevers as indicated. Black arrowheads indicate contact forces used in Fig. 2F. **(D-H)** HT1080 cells were treated by specific indicated siRNA’s (10 nM), or stably overexpressed with mock or lamin-specific vectors, and investigated either by AFS **(D-F)** or microaspiration **(G,H). (D)** Summary of elastic modulus measurements after indicated lamin modulation. Data points depict the normalized medians from elastic modulus measurements shown in Fig. S2 (with indicated P values), plotted against lamin expression values (from Fig. 1D). All data are depicted as ratio to normalized control (black point) on a double logarithmic (basis 2) scale. **(E)** Indentation at 20 nN compression force after depletion of indicated lamin isoforms. **(F)** Indentation at 20, 50 and 150 nN compression force after depletion of indicated lamin isoforms. **(G)** Representative image (from Video 2) of HT1080 cells treated with non-targeting control siRNA in the micropipette microfluidic device. Application of high pressure on one side of the device induces aspiration forces towards the respective other side of the device, with gradual pulling of cells and nuclei through 15 mm^2^ cross-sectioned channels over time. DNA was stained with Hoechst 33342. Image bar, 10 µm. **(H)** Left graph, averaged nuclear deformation length (± SEM; shaded area) over time after siRNA mediated knockdown. Significance was calculated after 8, 15, 30, 60 and 90 sec. Right graph, bars (mean) + SD after 120 sec. N = 3, 24-41 cells per condition. For all experiments, ***, P ≤ 0.001; **, P ≤ 0.01; *, P ≤ 0.05; ns, non significant. **(D-F)** Non-paired Mann-Whitney test, **(H)** two-tailed unpaired Students t-test.

**Figure 3.**
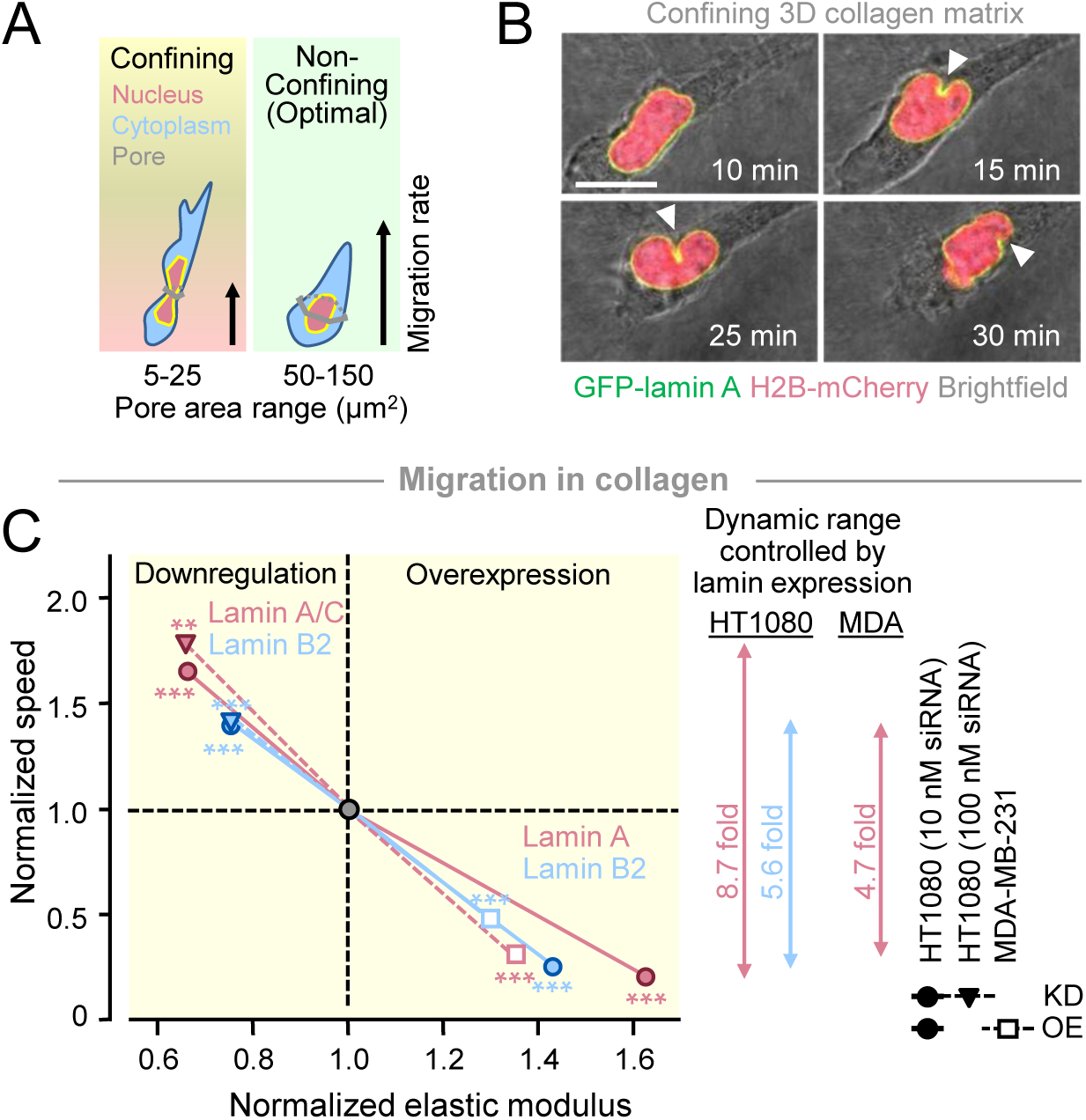
HT1080 tumor cell migration rates through dense collagen lattices inversely correlate with nuclear elasticity. **(A)** Scheme for classification of physical space into ‘confining’ or ‘non-confining’ (also ‘optimal’) pore areas relative to cell size, corresponding nuclear deformation, and migration capacity in collagen lattices. Whereas optimal pore sizes naturally match cross sections of proteolytically migrating cells and their ellyptic nuclei, confining pores induce dynamically adapting deformed nuclear shapes, leading to cell locomotion that is delayed by around half (visualized by arrow lengths; Wolf et al., 2013). Yellow/ red and green background coloring is used for the respective space conditions ‘confining/ very confining’ or ‘optimal’ in all following figures panels. **(B)** Sequence of cell expressing H2B-mCherry and GFP-lamin A migrating in confining collagen in the presence of GM6001 at indicated time steps (corresponding to Video 3). Arrowheads point to confined locations of the nucleus. Image bar, 10 µm. **(C)** Medians of migration rates from 3D confining collagen were plotted against medians of the elastic modulus measured from indicated cell lines after respective lamin modulation and as compared to normalized control (black circle). Values are from Figs. S2; S3E,F; and S4B,D. P values are indicated for migration rates, whereas non-significant elasticity values are underlaid by gray area. KD, knockdown; OE, overexpression.

As an independent assay, and to exclude potential confounding effects from apical actin fibers that span the nucleus in adherent cells and may thus affect the AFS measurements, we measured nuclear deformability of HT1080 cells in suspension using a microfluidic micropipette aspiration assay (Fig. 2G; Video 2). Cells depleted for lamin A/C and for lamin B2 showed similar, significantly increased nuclear deformation compared to non-target controls and lamin B1 depleted cells at short times indicative of nuclear stiffness (Fig. 2D, left). Over longer times, indicating viscous properties, nuclear deformation increased more in cells with downregulated lamin A/C than lamin B2 levels, with both cell types exhibiting significantly larger nuclear deformations than the NT and lamin B1 depleted cells. As consequence, after 120 seconds control cell nuclei became aspirated into the pipet to a length of 7.4 μm, yet aspiration was increased by 1.18-fold after lamin B2 knockdown and by 1.35-fold after lamin A/C downregulation (Fig. 2H, right).

To validate the relevance of lamin B2 in the control of nuclear mechanics, lamin expression was modulated in a number of additional cell types (Fig. S3A,D,G). AFS measurements confirmed, when compared to respective control values, a reduced stiffness after lamin B2 downregulation (0.66-fold) in MV3 melanoma cells whereas overexpression, vice versa, resulted in 1.30-fold increased elasticity in MDA-MB-231 breast cancer cells, respectively (Fig. S3B, E). Likewise, nuclear strain experiments in MCF10A breast cancer cells after lamin B2 overexpression showed a 1.5 fold reduced capacity of the cells to stretch,. In all measurements, these values were again exceeded by 0.62-fold after lamin A downregulation (Fig. S3B), and 1.35- and 4-fold after lamin A upregulation, respectively (Fig. S3E, H). In summary, downregulation of lamin A/C and lamin B2 increased, whereas lamin A/C and B2 overexpression reduced, the capacity of the nucleus to deform. In contrast, effects on lamin B1 in mediating stiffness to the cell nucleus were either not present (HT1080 cells; MDA-MB231 cells) or contrasted each other (MV3 and MCF-10A cells; Figs. 2D; S3B,E,H; Table 1). We therefore did not further follow up on a mechanical role of lamin B1 in mediating cell migration in confinement. Instead, we here conclude that lamin B2 follows lamin A/C in contributing to the viscoelastic behavior of the nucleus and determining its mechanical deformability.

### Lamin isoform-induced alterations in cell migration rates through dense collagen lattices

To assess whether the change in nuclear deformability associated with variation in lamin A/C and B2 levels affects migration efficacy in complex environments, HT1080 cells were tested in 3D fibrillar collagen lattices of different pore size ranges, based on previous quantitative pore calibration (Denais et al., 2016; Wolf et al., 2013). These included ECM lattices which either enable migration when nuclei deform (‘confining’; pore cross-section 5-25 μm^2^) or support fast migration without significant physical impact on nuclear shape (‘non-confining’; pores 50-150 μm^2^; generated by low collagen polymerization temperature or migration of cells in the presence of MMP-dependent collagen degradation; Figs. 3A, S4A). Accordingly, HT1080 cells moving through fibrillar collagen of confining density developed irregular shapes with local deformations (Fig. 3B), and with a speed of around 0.25 µm/min were below the speed range achieved in non-confining conditions (Fig. S4B-E; Video 3). After lamin A/C downregulation, migration speed in confinement increased to 1.65-1.8 fold, followed by a 1.2-1.55 fold increase after lamin B2 knockdown (Figs. 3C; S4B; Video 4). Notably, lamin isotype downregulation by 100 nM RNAi had a 10-30% stronger impact on migration, compared to 10 nM siRNA. Conversely, lamin overexpression caused a 4-5 fold speed delay after overexpression of lamins A, followed by B2 and lamin C (Figs. 3C; S4D; Video 5). Similarly, MDA-MB-231 cells in confining collagen conditions after downregulation of lamin A/C migrated 1.4-fold faster than non-targeted control cells (10 nM RNAi; data not shown), and upregulation of lamin A decreased migration speed to 30% as compared to control cell rates, followed by lamin B2 and C upregulation (Figs. 3C; S3F, left; Video 7). Notably, fast migration speed after lamin depletion was accompanied by decreased elastic modulus, and vice versa, for all lamin isoforms, though at somewhat different strength (Figs. 3C,D). Thus, by contributing to nuclear viscoelasticity, all lamin A/C and B2 mediate 5-9 fold different migration rates through confined spaces in fibrillar collagen.

Next, we examined whether changes in lamin levels affect cell migration speed in general, or only in confined conditions when the nucleus has to deform. In non-confining 3D collagen conditions migration depends on adhesion and cytoskeletal contractility-mediated mechanocoupling, but not on deformation of the nucleus (Fig. 3A) (Wolf et al., 2013). Here, similar cell migration rates in MDA-MB-231 and HT1080 cells were obtained, irrespective of lamin expression levels (Figs. S3F, right; Video 7; S4C,E). Likewise, when cells migrated across glass surfaces, median migration rates were not or only moderately decreased after up- or downregulation of lamin isoforms (Fig. S4F). These data indicate that lamin B2, like lamin A/C mediated migration speed in confining tissue space is controlled predominantly by nuclear mechanics rather than nucleo-cytoskeletal bonds.

### Altered cell migration rates through synthetic substrates of defined pore size after lamin deregulation

To confirm data in a spatially precise and complementary porous environment, we tested migration across transwell membranes with pores of uniform size and shape (either 5 or 8 µm diameter; Fig. 4A). Similar as to 3D collagen, transmigration of HT1080 cells through filter pores providing spatial confinement was increased after lamin downregulation and decreased after upregulation, with a similar contribution by each tested lamin isotype (Figs. 4B; S5A and B, each left). Transmigration data generated for MV3 melanoma cells showed a similar, 2-fold increase after lamin A downregulation, followed by lamin B2 (Figs. 4B; S3C). Generally, no changes were noted at non-confining pores (Figs. S3C, right; S5A and B, right), confirming that the basic migration ability of both cell types was unperturbed. As second engineered model, we used custom-developed microfluidic, polydimethylsiloxane (PDMS) migration devices that were functionalized with collagen and that provide precise confinement [Fig. 4C (Davidson et al., 2015)]. In addition, in contrast to polycarbonate transwell filters, these 3-dimensional environments are ideally suited for live cell imaging. The median transmigration rate of HT1080 cells through confining 10 µm^2^ constrictions was 15 minutes. Downregulation of lamins A/C accelerated the transit 1.7-fold, followed by lamin B2, whereas transit time through unconfined pores remained unaffected (Figs. 4D; S5C; Video 6). Together, the impact of the different lamin isoforms on migration efficacy in confinement is similar in both collagen models and synthetic substrates.

**Figure 4.**
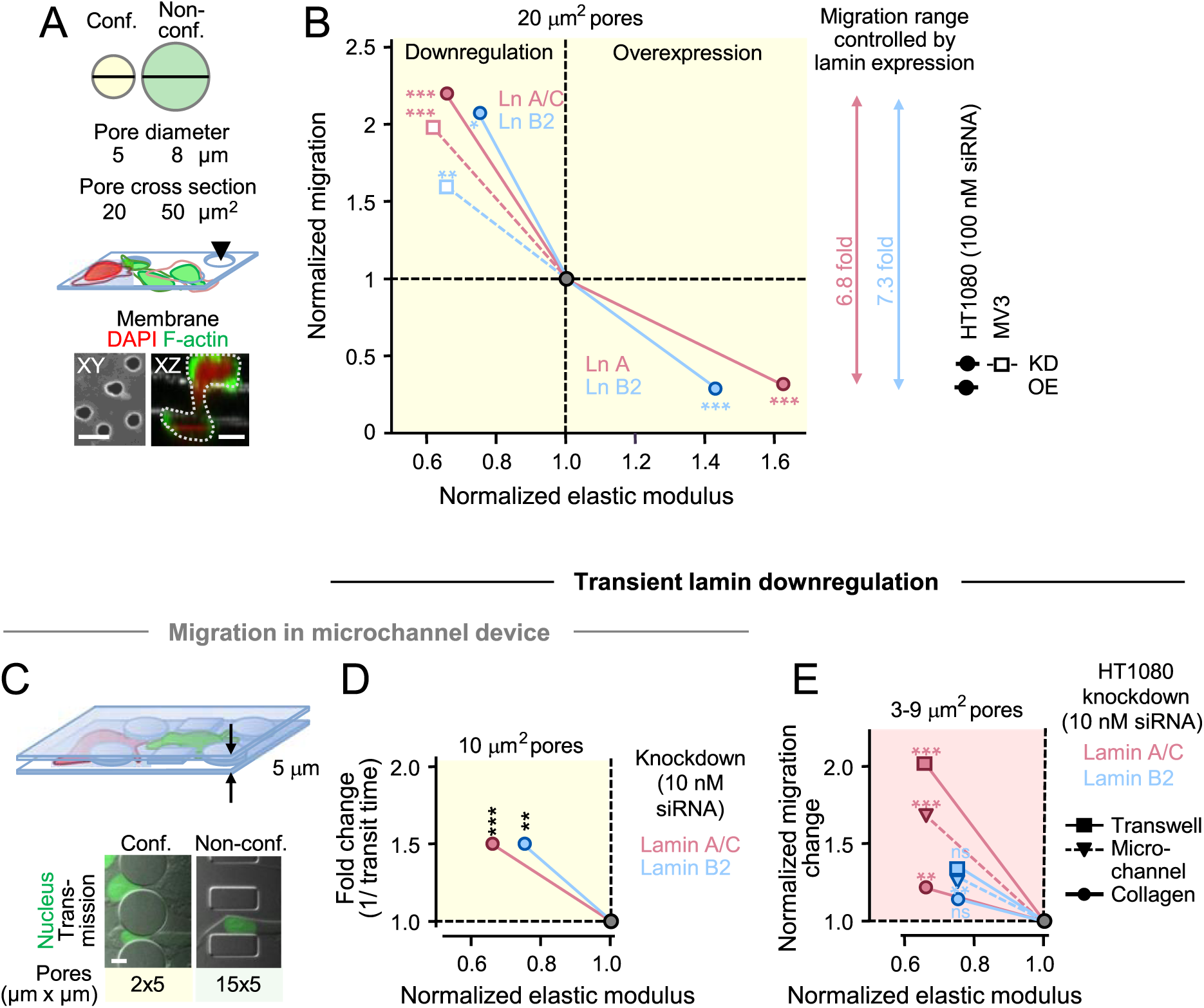
Migration rates in synthetic substrates of defined pore size. HT1080 cells were modulated for lamin expression as indicated and allowed to migrate through Boyden transwell chambers **(A-B)** or microchannels **(C-D). (A)** Scheme of used transwell pore sizes (upper panel) and transmigrating cells (middle). Conf., confining; Non-conf., non-confining. Bottom left, XY confocal depiction a transwell membrane containing 5 mm diameter pores; bottom right, XZ representation of deformed HT1080 cell (dotted outline) transmigrating a confining pore. **(B)** Migration rates through 5 mm diameter transwell pores plotted against elastic modulus measured from indicated cell lines after respective lamin modulation and compared to control (black circle). Values are the medians from Figs. S2; S3B,C; and S5A,B, and P values are indicated for transmigration rates. **(C,D)** Migration through microchannels. **(C)** Top, scheme depicting cells that migrate through the pores of the device; bottom, images of cells in the moment of transmigration through a 10 µm^2^ constriction or an unconstricted 75 µm^2^ channel (corresponding to Video 6). Nuclei are visualized through cell transfection with NLS-GFP. **(D)** Fold-decreased pore transit times through 2×5 mm diameter pores plotted against elastic modulus (dots depict the medians from Figs. S2A and S5C) measured after knockdown of indicated lamin isoforms as compared to control (black circle). P values are from transit times (here depicted as 1/ transit time resulting in increased values for faster pore transition). **(E)** Comparsion of HT1080 cell migration efficacies after downregulation of indicated lamin isoforms at near-limiting (3-9 mm2) pore conditions from the different migration models (depicted are the medians from Figs. S2A and S5E,F,G; also note MDA-MB-231 cell migration data through the microchannel device in Fig. S5H). P values are indicated for (trans)migration rates. ***, P <0.001; **, P <0.01; *, P <0.05; ns, non-significant). **(B,D,E)** All non-significant elasticity values are underlaid by gray area. All image bars, 10 µm.

To test whether a less rigid nucleus will allow migration at conditions that usually restrict migration due to limited nuclear deformation, cells were allowed to migrate within increased confinement (3-9 μm^2^ effective pore sizes; Fig. S5D). At these pore sizes, (trans)migration efficacy, as compared to confining conditions, further declined to 0.18 μm/min (collagen), 50% transmigration rate (transwell chamber) or 1.7-fold increased, 25 minute-long pore transit time (microchannel), indicating increasing mechanically impeding confinement (Fig. S5E-G). In most models, downregulation of lamin A/C significantly, and lamin B2 knockdown mildly, enhanced migration rates in HT1080 cells (Figs. 4E; S5E-G). Likewise, a 120 minute-long pore passage time by MDA-MB-231 control cells in a microdevice was accelerated after lamin A/C downregulation followed by lamin B2 in 5 μm^2^, but not in 75 μm^2^ pores (Fig. S5H). This nuclear deformation in strong confinement was accompanied by nuclear envelope (NE) rupture in 68% of all HT1080 cells, and was thus 40-fold increased when compared to non-confinement (Fig. S5I). NE ruptures further increased in small pores with depletion of each tested lamin isoform, in accordance with previously acquired MDA-MB-231 cell data (Denais et al., 2016). In summary, lamin B2 follows lamin A/C in regulating migration efficacy in confining and near-limiting 3D, but not in unconfining 3D or 2D environments.

### Lamin expression-dependent nuclear deformation during migration

The data so far indicated that lamin B2, like lmin A/C, modulates cell speed primarily through regulating the viscoelasticity of the nucleus, and thereby facilitates or impede migration in mechanically demanding 3D environments. We therefore tested whether lamin regulation directly affects the maximal deformation of nuclei during 3D migration. As a control, downregulation of lamin isoforms yielded near-equal nuclear diameters between around 10 - 11 μm.in yet round HT1080 cells suspended into a 3D network in the moment of collagen polymerization (Fig. 5A). After 15-20 hours of migration, when control cells developed nuclear diameters of 7 μm, lamin downregulation supported 120-140% increased deformations (Fig. 5B,C). The percentage of nuclei where the smallest nuclear diameter was ≤3 µm, corresponding to the ‘physical migration limit’ (Wolf et al., 2013), increased from 3% in control cells to 8-15% in cells where lamins were downregulated, with notable shape pleomorphism ranging from elliptic to multi-indented shapes (Fig. 5B). To address whether lamin expression regulation exceeds the limit of nuclear deformation during migration, only the diameters of deformed nuclei were measured. Lamin downregulation decreased the smallest nuclear diameters of 4.2 µm in control cells down to 3 µm, and reciprocally increased the number of nuclei with extreme deformation from 7% in control cells to 23% for lamin B2 and to 50% for lamin A/C (Fig. 5D). Vice versa, lamin upregulation in HT1080 cells led to 1.2 to 1.5-fold enlarged cytoplasmic protrusions in confining collagen (Fig. 6A, upper row; B; Video 5), as well as to rounded nuclei reminiscent of migration-abrogated cells (Figs. 6A, bottom row; S5D) (Wolf et al., 2013). Nuclear diameters of 6.5 μm from control cells increased by 26% after lamin B2, and more dominantly (by 45%) after lamin A overexpression in HT1080 cells (Fig. 6C). Similarly, an increase of nuclear diameters from around 4 to 6 μm (150%) in MDA-MB-231 cells occured in dense, but not in loose, matrix (Figs. 6D,E). Accordingly, not more than 2% cells per population reached nuclear diameters of ≤3 μm (Fig. 6C,E), in contrast to numbers after lamin downregulation. Together, these data indicate that excess expression of lamins A or B2 decreases the capacity of the nucleus to deform, thereby exhausting migration at pore sizes that are usually confining. Thus, nuclear diameters during migration are controlled by lamin isoforms A/C and B2, concordant to the cells nuclear mechanics and inverse to migration speed (Fig. 6F,G).

**Figure 5.**
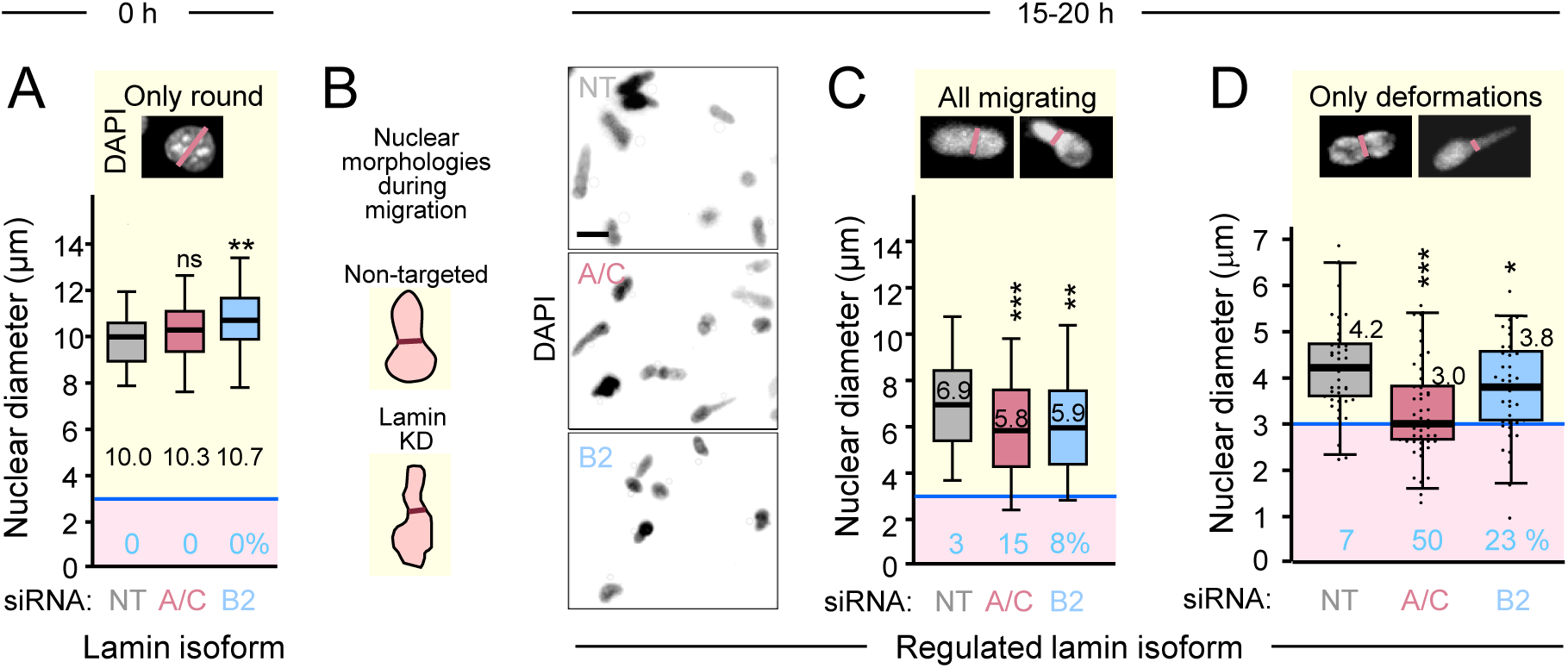
Reduced nuclear diameters after lamin downregulation during HT1080 cell migration in dense collagen. After treatment with non-targeting or lamin-specific siRNA, all cells were cultured in collagen (rat tail, 1.7 mg/ml) in the presence of GM6001. **(A)** Diameters of individual cell nuclei, measured from DAPI signal as indicated by the red lines in the example images (top panels) immediately after collagen polymerization. **(B)** Left, scheme of typical nuclear shapes developing during migration in confinement; red lines, ‘minimal’ nuclear diameters. KD, knockdown. Right, representative images of nuclear morphologies after 15-20 h migration. Lattices were fixed, stained, and imaged for DAPI (together with phalloidin fluorescence and transmission signal for intact cell morphology, not shown), and confocal z-scans were maximally projected. Image bars, 10 mm. **(C,D)** Diameters of individual cell nuclei.after 15-20 h migration. **(A,C,D)** Horizontal lines, boxes and whiskers show the medians, 25^th^/75^th^, and 5^th^/95^th^ percentile. ***, P <0.001; **, P <0.01; *, P <0.05; ns, non-significant (non-paired Mann-Whitney test). Red areas with blue horizontal line in the lower part of the graphs, and numbers (in %) depict the range and percentage of nuclei at or below 3 mm diameter. Black numbers in graphs indicate median nuclear diameters. **(A)** N = 2-3; 48-81 cells; **(C)** n = 2; 80-120 cells; **(D)** n = 3; 36-46 cells per condition.

**Figure 6.**
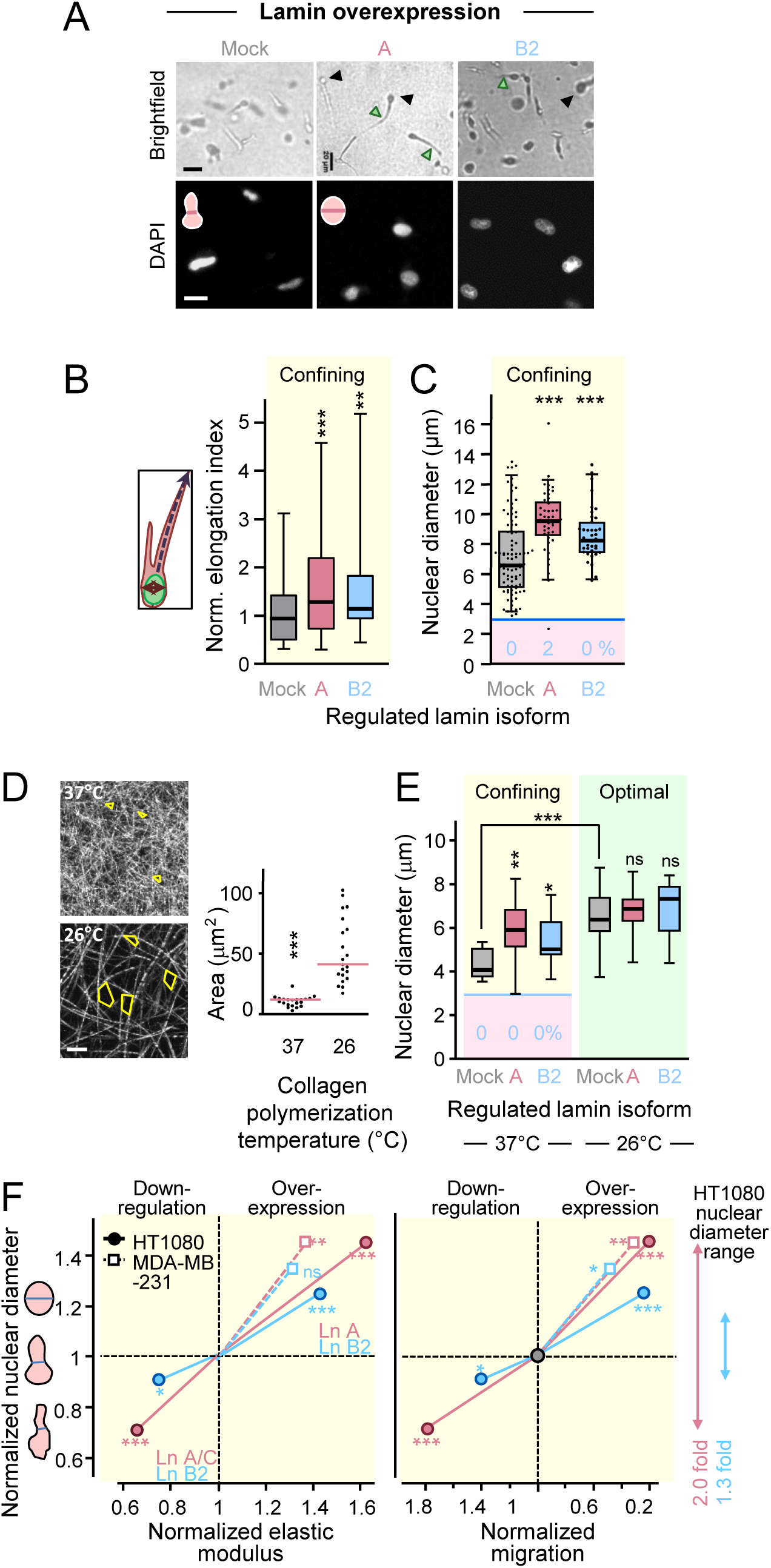
Elongated cell shapes and rounded nuclei during migration in dense collagen after lamin upregulation. **(A-C)** HT1080 or **(E)** MDA-MB-231 cells were stably transduced with control or indicated lamin vectors and migrated within rat tail **(A-C)** or bovine **(E)** collagen (1.7 mg/ml; unless indicated otherwise) in the presence of GM6001 for 15-20 h. **(A)** Top row, representative brightfield images of cells with indicated lamin modulations after migration in collagen for 10 h (corresponding to Video 5). Black arrowheads, round cell rears indicative of nuclear position; green arroheads, elongated cell protrusions. Bottom row, representative maximum projected DAPI images from nuclei of migrating cells. **(B)** Cell elongation. Left, schematic depiction of calculated elongation index measured as cell body length divided by cell body width. Right, quantified normalized nuclear elongation index from cells as shown in A). N = 2-4; 41-125 cells per condition. **(C)** Quantification of minimum nuclear diameters as shown in A. N = 1-6; 24-90 cell nuclei per condition, shown as dots. **(D)** Left, bovine collagen lattices of 2.5 and 1.7 mg/ml, and at 37°C and 26°C polymerization temperatures, respectively, imaged by confocal reflection microscopy, with depiction of pore areas (yellow outline). Right, pore area quantification. **(E)** Minimum nuclear diameters after MDA-MB-231 migration for 20 h in collagen polymerized at indicated temperatures. 8-26 cells per condition. **(F)** Correlation of normalized nuclear diameters in confinement plotted against elastic modulus (left) or migration rates (right). Dots are the medians from Figs. 5F; 6C; S2; S4B,D (HT1080); 6E and S3E,F (MDA-MB-231). P values are from the calculation of nuclear diameters, and all non-significant elasticity values are underlaid by gray area. The nuclear deformation range during migration by experimental lamin modulation is indicated on the right. **(C,E)** Red areas with blue horizontal lines in the lower parts of the graphs mark ≤3 mm nuclear diameter ranges. Numbers in blue, percentages of cell nuclei below ≤3 mm. **(B,C,E)** Horizontal lines, boxes and whiskers show the medians, 25^th^/75^th^, and 5^th^/95^th^ percentile. ***, P <0.001; **, P <0.01; *, P <0.05; ns, non-significant (non-paired Mann-Whitney test). Image bars, 20 μm (**A**, upper row); 10 μm (**A**, lower row; **D**).

### Increased dynamic nuclear shape change after lamin downregulation

Lastly, to examine whether lamin-mediated nuclear deformation underlies nuclear shape dynamics, we analyzed the morphological changes that cell nuclei undergo during constricted migration. Sequences from in-focus nuclei during cell migration were collected and nuclear morphologies extracted (Fig. 7A, Video 8). Subsequent nuclear shapes were rotated for a maximal overlap to take directional changes during migration into account and remaining changes between irregular shapes over time were calculated as nuclear shape fluctuation (Fig. 7B) (Krause et al., 2019). These shape changes increased most after downregulation of lamin A/C and were again followed by lamin B2 downregulation (Fig. 7C; see Video 8). Thus, these lamins are important regulators of shape maintenance in a graded fashion, where depletion of each lamin isoform increases nuclear deformability and shape adaptation for increased migration rates in confinement with heterogeneously formed pores (Fig. 7D). In summary, by regulating nuclear mechanics, A/C and B2 alone or together control nuclear deformation for migration rates and nuclear integrity.

**Figure 7.**
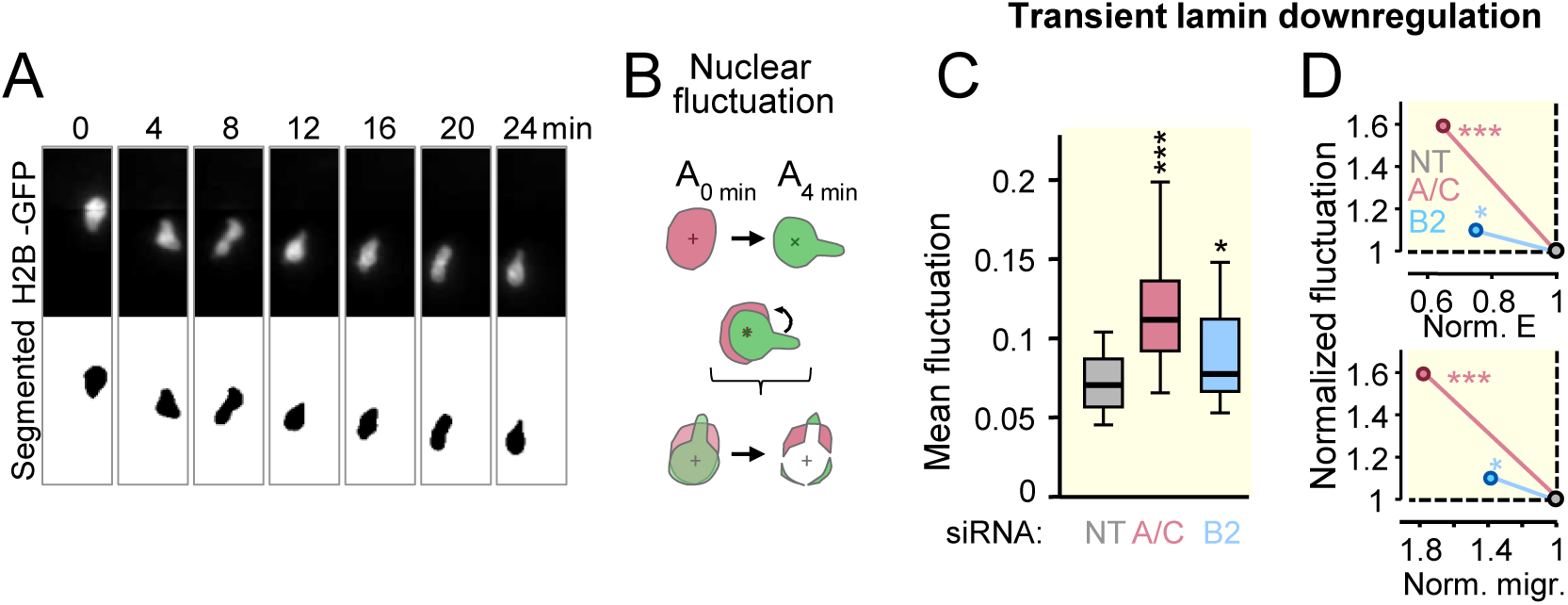
Increased nuclear shape change during migration in small spaces after lamin downregulation. HT1080 cells stably expressing H2B-GFP and migrating at confining conditions (rat tail collagen; 1.7 mg/ml, in the presence of GM6001) were imaged by dynamic spinning disk microscopy for changes in nuclear morphology and analyzed. **(A)** Representative image sequence of a nucleus during cell movement (upper row), that remained in focus and was thresholded and segmented for further analysis (bottom row; corresponding to Video 9). **(B)** Depiction of analysis principle of nuclear fluctuation. Nuclear areas A from 2 consecutive time points, having developed different shapes (upper row), were overlaid at the center (cross) and rotated for maximal overlap (middle row; indicated by bended arrow) to remove errors by changes in step angle during the migration process from analysis. The lack of overlap was calculated as remaining pixel difference (colored areas) between two time points. **(C)** Mean values of the fluctuations from each nuclear shape to the next per cell migration sequence after indicated control and silamin treatments, calculated as stated in method section. A fluctuation value of 0 is equal with a complete overlap, and a (theoretical) value of 1 means no overlap at all. N = 3-4; 18-32 cells per condition. Horizontal lines, medians; box plots, 25^th^ to 75^th^ percentile; and whiskers, 5^th^ to 95^th^ percentile. For all experiments, non-paired Mann-Whitney test was used for statistical analysis. ***, P <0.001; **, P <0.01; *, P <0.05; ns, non-significant. **(D)** Fluctuation plotted against elasticity and migration after depletion of indicated lamin isoforms. Dots depict the medians from Figs. S2A, S4B and 7C, and P values are from Fig. 7C.

## Discussion

By probing migrating tumor cells with low or high lamin isoform expression levels in 3D substrate of optimal, confining or near-limiting porosity relative to cell size, we have identified roles for each lamin isoform in nuclear mechanics and migration. Besides A-type lamins (Davidson et al., 2014; Harada et al., 2014), we show that the lamin isoform B2, but not B1, regulates nuclear material properties, including elasticity, viscoelasticity and deformation in a non-redundant manner. Perturbation of lamins A/C and B2 caused changes in nuclear deformation during constricted migration. These ranged from complex shape adaptations prone to nuclear envelope rupture after lamin downregulation to a poorly deformable, roundish nucleus after lamin overexpression. Consequently, the reduction of each lamin isoform increased cell migration, whereas upregulation caused strong migration delay (Wolf et al., 2013), thus regulating speed by up to 9-fold (Fig. 8). Our data may imply the existence of a cooperating lamin network, with all lamin isoforms fulfilling non-redundant as well as overlapping functions in regulating nuclear shape adaptation and integrity during migration in 3D environments.

**Figure 8.**
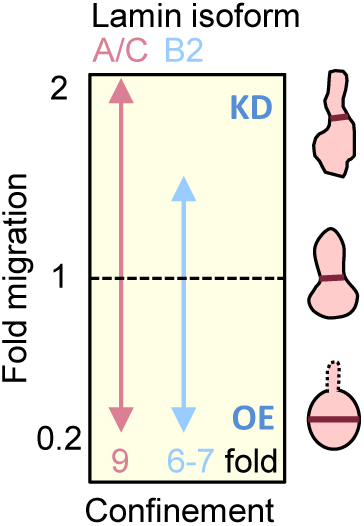
Lamin A/C and B2 expression determines nuclear deformation and cell migration efficacy in dense 3D environments. Left, summary on the impact of lamin expression regulation on relative migration efficacy at confining pore size. Migration of cells not modulated for lamin expression is set to 1. Right, respective nuclear shapes and diameters (brown line) concordant to lamin expression level during migration in confinement. KD, lamin knockdown; OE, lamin overexpression.

### Regulation of nuclear mechanics and cell migration by B-type lamins

According to their early presence of in soft proliferating stem cells and lack of regulation by surrounding tissue stiffness (Swift et al., 2013), B-type lamins have been accounted a role in nuclear organization, rather than contributing to nuclear mechanics. This view was supported by the lack of lamin B1-mediated effects in nuclear deformation measurements under strain, using *Lmnb1*^Δ/Δ^ mouse embryonic fibroblasts (Lammerding et al., 2006). Others reported that nuclear stiffness as measured by AFS increased with lamin B1 expression in skin fibroblasts after *LMNB1* gene duplication in autosomal dominant leukodystrophy patients, as well as after ectopic lamin B1 expression in HEK293 and neuronal N2a cells (Ferrera et al., 2014). On the contrary, lamin B1 depletion stiffened the nucleus as shown by micropipette aspiration (Shin et al., 2013), especially when lamin A/C levels were low (Stephens et al., 2017), as well as increased nucleolar stiffness as shown by AFS (Louvet et al., 2014). These contradicting results were consistent with our own results. Thus, overall, the impact of lamin B1 on the maintenance of cell stiffness may be cell type-dependent, but may also depend on the levels of other lamin isoforms.

Because of the overall not apparent or contrary effects mediated by especially lamin B1, a contribution of lamin B2 to nuclear mechanics was overlooked so far. Here, by combining AFS and micropipette aspiration to mechanically deform the nucleus to similar extents as nuclear deformations required for 3D migration, we found a robust contribution of lamin B2 to nuclear mechanics. Following lamin A/C in magnitude, the contribution of lamin B2 to nuclear deformability was revealed in four independent cell models and by three different techniques, and therefore likely can be generalized.

### Cooperating lamin networks

The impact of lamin B2 levels on nuclear deformability and migration implies non-redundant and overlapping functions with A-type lamins in stabilizing nuclear mechanics. Indeed, originally thought to form separate layers (Goldberg et al., 2008a; Goldberg et al., 2008b), A- and B-type lamins form strongly interlinked networks of particular porosity (‘face areas’) to regulate stability of the nucleus in MEF cells (Shimi et al., 2015; Turgay et al., 2017). Accordingly, knockout of the *LMNA* gene mildly increased the porosity of the B-type lamin network (Shimi et al., 2015). This effect might be supported by a biochemical study demonstrating that the stability of the lamin B2 network requires interaction with the other, i.e. A-type lamin polymer network (Schirmer and Gerace, 2004). Together, this may show that the structural meshwork properties of individual lamins depend on the presence of the respective other lamin isotypes.

### Implications for cell navigation in complex environments

To rule out a major effect on the general migration machinery after lamin depletion (Corne et al., 2016; Hale et al., 2008), we tested migration across ECM-coated glass and large pore 3D conditions and detected no principal migration defect when cells move in non-confining conditions. When, however, comparing speed changes under confinement, downregulation of lamins allow acceleration by 1.5-fold in 3D collagen and PDMS channels and up to 2.5-fold crossing transwell membranes. Conversely, upregulation of all lamin isoforms resulted in around 20-25% remaining migration rates in both 3D collagen or transwell membranes. This indicates that adaptation of nuclear deformation is the primary mechanism to facilitate or impede cell migration in 3D confinement. Thus, irrespective of cell type, ligand density, 3D substrate stiffness, or the (ir)regularity of pore shapes, lamin expression levels control nuclear deformability and migration rates. By regulating the ability of cells to move into and through dense tissue, with a total amplitude of 870% from fastest to slowest condition, lamin expression thus represents a central regulator of cell mechanics for cell navigation through 3D tissue. Cells with high lamin levels and limited deformability may retain full ability to navigate through ECM environments of wide space, including linear tracks present in the connective tissue and between myofibers in the living organism. In addition, cells with stiffer nuclei may retain a stronger ability to deform and compress the tissue during migration, which may support proteolytic remodeling and tissue disruption (Wolf et al., 2007). Conversily, cells with low lamin expression may additionally be able to disseminate through dense and complex-shaped ECM, including irregular clefts between adipocytes or traverse across dense basement membranes (Glentis et al., 2017; Weigelin, 2012). Thus, rather than providing a stop signal for migration, nuclear deformability may dictate routes of preference based on available space and alignment of tissue structures.

### Implications for cancer progression

Besides regulating cell migration, A-and B-type lamins may cooperate in protecting the cell nucleus against mechanical assault caused by deformation and extracellular shear stress. When being deformed towards high nuclear curvature by moving individually through dense

ECM, cancer cells and leukocytes with low lamin A/C, B1 or B2 expression all experience nuclear envelope ruptures, followed by DNA damage and, potentially, genomic instability, nuclear fragmentation and cell survival (Denais et al., 2016 ; Harada et al., 2014; Hatch et al., 2013; Hatch, 2016; Irianto et al., 2017; Raab et al., 2016; Xia et al., 2018). Thus, downregulation of A- and B-type lamins may impose a dual effect on cancer progression, by facilitating dissemination into a vast range of tissue spaces, as well as supporting invasion-associated DNA damage and, potentially, cancer aggressiveness. Conversely, lamin upregulation may mechanically protect the nucleus by limiting deformation and steering migration into more defined tissue routes. Because intratumor heterogeneity with both up- and downregulated lamin expression is a typical feature of neoplasia (Chow et al., 2012; de Las Heras et al., 2013), it remains to be investigated whether and how invasion patterns and nuclear integrity status including DNA damage and cell survival are affected by lamin deregulation during cancer disease.

## Abbreviations

2D: two-dimensional;
3D: three-dimensional;
AFS: atomic force spectroscopy;
ECM: extracellular matrix;
H2B: histone-2B;
MMP: matrix metalloproteinase;
NII: nuclear irregularity index;
NT: non-targeting control;
PDMS: polydimethylsiloxane

## Acknowledgements

We thank Celine Denais and Kathy Zhang for help with the microfluidic migration device assays; Jos Broers for providing the GFP-laminA vector; Esther Wagena and Stephanie Alexander for the generation of stable cell lines and western blotting; Lisette Meerstein and Louis Wolf for help with image processing and analysis; Ineke van der Zee for help with statistical analysis; Ben Fabry for helpful discussions on AFS analysis; the RIMLS Microscopic Imaging Center for the use of facilities, in particular the combined AFS-confocal microscope (NWO Medium Sized Investment Grant 91110007); as well as the Cornell NanoScale Facility (CNF) where microdevices were fabricated (NNCI member supported by NSF Grant NNCI-1542081).

This project was supported by the Netherlands Science Organization (NWO-VENI 680.47.421 to JtR; NWO-VIDI 917.10.364 to K.W.; NWO-VICI 918.11.626 to P.F.) and the Dutch Cancer Foundation KWF (grant number 11199 to K.W); by the Leverhulme Trust Research Project Grant (RPG-2014-149 to FWY and AM) and the European Union’s Horizon 2020 research innovation program (Marie Sklodowska-Curie grant 642866 to AM); and by National Institutes of Health awards (R01 HL082792 and U54 CA210184 to JL); a Department of Defense Breast Cancer Breakthrough Award (BC150580 to JL) and a National Science Foundation CAREER award (CBET-1254846 to JL).

## Competing interests

The authors declare no competing financial interests.

## Separate author contribution

K.W. and P.F. conceived and designed the experiments and were supported by J.L. and J.tR. M.VK., M.KtL., J.tR., V.tB., R.M., R.P., P.I., T.vO., M.Z. and K.W. performed the experiments. M.VK., M.KtL., J.tR., V.tB., R.M., R.P., P.I., T.vO., F.Y., M.S. and K.W. analysed the data. J.L. and M.Z. contributed material, and J.tR., A.M. and F.Y. contributed analysis tools. K.W., P.F. and M.VK. wrote the paper.

All authors read and corrected the manuscript.

## Materials and Methods

### Antibodies and dyes

The following antibodies, isotypic IgGs and fluorescent dyes were used: monoclonal mouse anti-lamin A/C IgG1 (clone 131C3; Acris), polyclonal goat anti-lamin A/C (sc6215; Santa Cruz Biotechnology), polyclonal goat anti-lamin B1 (sc6217-M20; Santa Cruz Biotechnology), monoclonal mouse anti-lamin B2 IgG (clone X223; Progen), polyclonal rabbit anti-lamin B2 (10895-1-AP; ProteinTechGroup); monoclonal mouse anti-β-tubulin IgG1 (clone E7; Developmental Studies Hybridoma Bank, University of Iowa); monoclonal rabbit anti-GAPDH IgG (clone 14C10; Cell Signaling Technology) and polyclonal anti-β-actin (Abcam) were used for confocal microscopy or western blotting. Secondary goat anti-mouse IgG and rabbit anti-goat IgG conjugated to horseradish peroxide (ECL; Thermo Fisher Scientific), or a fluorophore specific to the Odyssee system (IRDye 680RD and IRDye 800RD; LI-COR Biosciences) were used for western blotting, and secondary Alexa Fluor 488-conjugated pre-absorbed goat anti-mouse, donkey anti-goat and goat anti-rabbit IgGs (Invitrogen), Alexa Fluor 488 or 647-conjugated phalloidin (Invitrogen), and DAPI (Sigma) were used for confocal microscopy.

### Cell lines and lamin expression modulation approaches

The following cell lines were used: human HT1080 fibrosarcoma cells (ACC315, DSMZ Braunschweig, Germany), MV3 melanoma cells (Maaser et al., 1999), MDA-MB-231 breast cancer cells (Moss et al., 2009) and MCF10A breast epithelial cells. HT1080, MV3 and MDA-MB-231 cells were cultured at 37°C and 5% CO_2_ humidified atmosphere in Dulbecco’s modified Eagle medium (DMEM; Invitrogen), supplemented with 10% fetal calf serum (FCS; Sigma Aldrich), 2 mM L-glutamine, 1 mM sodium pyruvate (Invitrogen) and 100 U/ml penicillin and 100 μg/ml streptomycin (PAA). MCF10A cells were cultured in DMEM/HAM12 medium supplemented with 5% horse serum, 0.5 µg/ml hydrocortisone, 5 µg/ml insulin, and 10ng/ml EGF. Cell lines were used untreated; or modified for either lamin expression as described below, or for stable expression of fluorescent markers such as nuclear histone-2B (H2B)-coupled GFP, H2B-mCherry, NLS-GFP, or GFP-lamin A, alone or in combination (Denais et al., 2016). Briefly, for the generation of HT1080 cells stably expressing GFP-Lamin A, lamin A DNA in a pEGF-C1 vector (provided by J. Broers, Maastricht University) was transfected into wild type HT1080 cells by Lipofectamin 2000, selected with neomycin for stable expression, sorted twice, and clonally selected for moderate GFP-lamin A expression [clone 1, (Denais et al., 2016)]. Further, after transduction with lentiviral nuclear H2B-coupled GFP or H2B-mCherry constructs, HT1080 cells were selected by blasticidine and FACS sorted for high expression. To modulate lamin expression, cells were either treated with specific siRNAs for transient lamin knockdown or stably transduced with vectors encoding lamin genes.

For transient lamin knockdown, cells were cultured in antibiotics-free supplemented DMEM in 6-well plates (each 250.000 cells) for 24 h. Cells were treated with a pool of small interfering (si) RNAs consisting of 4 single RNAs each and targeting expression of lamin A/C, lamin B1, or lamin B2 protein, or non-targeting (NT) negative control (on-target plus, SMARTpool; Dharmacon). The forward 21-nucleotide siRNA sequences for the NT control were 5-UGGUUUACAUGUCGACUAA-3, 5-UGGUUUACAUGUUGUGUGA-3, 5-UGGUUUACAUGUUUUCUGA-3, 5-UGGUUUACAUGUUUUCCUA-3; for si*LMNA* the forward sequences were 5-GAAGGAGGGUGACCUGAUA-3, 5-UCACAGCACGCACGCACUA-3, 5-UGAAAGCGCGCAAUACCAA-3, 5-CGUGUGCGCUCGCUGGAAA-3; for si*LMNB1*, the forward sequences were 5-AAUAGAAGCUGUGCAAUUA-3, 5-GAUCAAGCUUCGAGAAUAU-3, 5-GCAUGAAACGCGCUUGGUA-3, 5-GCGCAAGCCCUUCAUGAGA-3 and for si*LMNB2* the forwards sequences were 5-UAACGCGGAUGGCGAGGAA-3, 5-GAGAUCGCCUACAAGUUCA-3, 5-GUGAAGAAGUCCUGGUGA-3, 5-CCUCGACGCUGGUGUGGAA-3. siRNAs were transferred into cells with Dharmafect 4 transfection reagent according to the manufacturer’s protocol and cultured with antibiotics-free DMEM for 48 h prior to characterization and functional studies. Viability of cells after exposure to Dharmafect reagent and specific RNAi’s was confirmed by the lack of membrane damage detected by staining with propidium iodide (Sigma) and flow cytometry, which resulted in 91-97% propidium iodide-negative cells at each condition, indicating membrane integrity; and intact migration function (see Figs. S4B, C for lamin-specific siRNA during migration compared to untreated control cells, or within collagen providing optimized pore conditions).

For stable lamin overexpression in HT1080 wildtype cells, human lamin *LMNA, LMNC, LMNB1 or LMNB2* DNAs [UniProt P02545-1 (prelamin A); P02545-2 (lamin C); P20700 (lamin B1); and Q03252 (lamin B2 short version according to (Schumacher et al., 2006)] were cloned into lentiviral PCDH-EF1-MCS1-puro vectors (System Biosciences).

Lentiviruses were produced by cotransfection of lentiviral target plasmids (ENV, Pol, GAG) with the respective lamin containing or empty (Mock) vectors into HEK 293-TN cells using PureFection nanotechnology-based transfection reagent (System Biosciences). The medium was exchanged the next day and replaced by antibiotics-free DMEM containing 10% FCS. Lentivirus-containing supernatants were collected at 48 h and 72 h after transfection, filtered through a 0.45 µm filter and used as viral stock that HT1080 cells were transduced with for at least 3 consecutive days, in the presence of 8 µg/mL freshly prepared polybrene (Sigma-Aldrich). The viral solution was removed and cells were incubated in fresh medium for further 24 h before then being sub-cultured under continuous puromycin selection. For the production of retrovirally transduced MDA-MB-231 cells, retroviral target plasmids (ENV, Pol, GAG) were co-transfected with the respective lamin-transfected pRetroX-IRES-ZsGreen vectors into HEK 293 cells and the procedure proceeded as for the lentivirally transduced HT1080 cells. Retrovirally transduced MDA-MB-231 cell lines were tested by reverse transcriptase (PERT) assay (University of Zürich, Switzerland), and the absence of retrovirus production and secretion into the cell culture supernatant was confirmed. Because of the absence of a neomycin resistance gene in retrovirally transduced MDA-MB-231 lines, cells were FACS-sorted and the stability of lamin expression confirmed by flow cytometry for over 10-15 passages, during which experiments were performed.

### SDS-Page and Western Blotting

Lamin knockdown and overexpression efficiencies were determined by electrophoresis and western blot analysis from whole-cell lysates (62.5 mM Tris-HCl; 2% w/v SDS; 10% glycerol; 50 mM DTT; 0.01% w/v bromophenol blue), followed by either chemiluminescence detection (ECL detection kit; GE Healthcare) or fluorescence detection (Odyssee; LI-COR Biosciences) and densitometric analysis (Fiji ImageJ).

### Nuclear deformability measurements by AFS

Imaging and quantification of cell deformation, elastic modulus and dissipation were performed on a Catalyst BioScope atomic force microscope (Bruker, Santa Barbara, CA) combined with a 3-channel confocal microscope TCS SP5 II (Leica; 20×/NA 0.70 or 40×/NA 0.85 air objectives). The measurements on nuclear deformation and elastic properties, including the conversion of force-deformation into force-indentation (F-δ) curves were performed as described (Krause et al., 2013). Briefly, 1 or 2 days before experiments, 40,000 or 80,000 cells were seeded in a Willco dish supplemented with DMEM/10 % FCS. Cells were probed by defined forces and at a speed of 10 μm/sec using calibrated cantilevers coupled to 10 μm-diameter beads (NP-S, type D, nominal spring constant of 0.06 N/m, Bruker, for the calculation of the elastic modulus; or NSG11, type B, nominal spring constant of 5.5 N/m, NT-MDT, for the generation of the approach-retraction sequence; see Fig. S2A). After the measurements and after baseline correction, F–δ curves were fitted over a 0-up to 1000 nN range. To calculate the elastic modulus, an in-house written Igor Pro 6 (WaveMetrics) algorithm was applied using the simplified Hertz model for spheres in contact with a flat surface and deformations less than 10% of the original height, thus without the requirement to correct for sample affinity and non-linearity (Krause et al., 2013; Lin et al., 2009).

For the visualization of cell deformation by AFS probing optically assisted AFS measurements were performed with a speed of 250 nm/s and using fluorescent Crimson-labeled 10 µm-diameter beads (Invitrogen), as previously published (Krause et al., 2013).

### Micropipette aspiration assay

PDMS microfluidic devices were fabricated and used as previously described (Denais et al., 2016). In brief, the device was passivated with 2% BSA and 0.2% FCS blocking solution to prevent cell adhesion to the PDMS and cells were subsequently perfused into the device. The device was pressurized at 1-2 psi on the cell side to deform the nuclei into the constrictions. Dimensions of the micropipette channels were 5 µm in height and 3 µm in width. Imaging was performed at 37°C on an inverted Zeiss Observer Z1 microscope equipped with a CCD camera (Photometrics CoolSNAP KINO) using a 20×/NA 0.8 air objective, and image acquisition was automated through ZEN (Zeiss) software with imaging intervals of 2 sec collecting images for DIC and GFP. Nuclear transit times were determined by a custom-written automated image analysis program as described previously (Elacqua et al., 2018).

### Nuclear strain experiments

Substrate strain experiments to measure nuclear stiffness were performed as described previously (Lombardi et al., 2011; Zwerger et al., 2013). In brief, cells were plated on fibronectin-coated silicone membranes, incubated with Hoechst 33342 to DNA, and subjected to ∼20% uniaxial strain using a custom-build microscope-mounted strain device. Nuclear strain was calculated based on fluorescence images of the cell nucleus before, during, and after strain application and normalized to the applied membrane strain. Cells that became damaged or detached during the stain application were excluded from the subsequent analysis.

### Preparation of 3D collagen lattices

Acidic rat-tail collagen type I solution [Becton Dickinson, BD, for all experiments unless indicated otherwise; or from S. Weiss lab (Wolf et al., 2013), for Fig. S4A only]; or acidic bovine collagen type I solution (Nutragon, Advanced BioMatrix; for all experiments with MDA-MB-231 cells) were used to generate fibrillar collagen matrices (1.7 or 2.5 mg/mL) for 3D migration experiments. Lattices were reconstituted in vitro by raising the pH to 7.4 using either NaOH and buffering by 25 μM Hepes (Sigma-Aldrich) for rat tail collagen, or 0.75 % Na-bicarbonate solution (Gibco) for bovine dermal collagen, together with minimum essential Eagle’s medium (Sigma-Aldrich), and were polymerized at 37°C unless indicated otherwise. Collagen lattices were used either cell-free or with cells (20.000/100 μl), which were suspended in neutralized collagen solution before polymerization and placed as droplet into a well insert or self-constructed chamber (Wolf et al., 2013). After polymerization, supplemented medium was added and, where indicated, broad spectrum matrix metalloproteinase (MMP) inhibitor GM6001 (Ilomastat; 20 μM, unless stated otherwise; Calbiochem) was added to both collagen-cell suspensions and supernatants.

### Time-lapse-microscopy and quantitative cell tracking

Cells seeded at low density on glass bottom Willco dishes or cell-collagen cultures were monitored by temperature- and CO_2_-controlled bright-field inverse microscopy (37°C, 5%; air objectives, 10×, NA 0.20, Leica; 20×, NA 0.30, Leica; and 10×, NA 0.25, Nikon DiaPhot 300). Images were collected by CCD cameras (Sentech or Hamamatsu C8484-05G), and either the 16-channel frame grabber software (Vistek) or the 2D time lapse software version 2.7 (Okolab) was used at 4-minute frame intervals for up to 30 h (Wolf et al., 2013). Migration speed was quantified by computer-assisted cell tracking [Autozell 1.0 software; Center for Computing and Communication Technologies (TZI), University of Bremen, Germany] for the generation of XY paths with 4-12 min step intervals. The average speed per cell was calculated from the length of the path divided by time, including “go” and “stop” phases. Only viable cells that showed dynamic changes in cytoplasmic extensions and retractions including mitosis were tracked and included into the analysis.

### Boyden chamber transmigration assay

Transwell migration experiments were performed as described previously in (Wolf et al., 2013). Briefly, 6.5 mm diameter transwell chambers with uncoated polycarbonate filters of 8, 5, and 3 μm diameters pore size (BD and Costar) were filled with 30.000 cells suspended in DMEM supplemented with 2% FCS. Cells migrated through the porous membrane into the lower transwell chamber containing supplemented DMEM/ 10% FCS and pro-migratory agent lysophosphatidic acid (LPA; 10 μM; Sigma) for 20 h. The remaining cells on the upper membrane side were removed by a cotton swab, and the transmigrated cells attached to the lower membrane side were fixed, stained with crystal violet, embedded by mowiol, imaged and counted.

### Microfluidic migration devices

PDMS microfluidic devices were fabricated and used as previously described (Davidson et al., 2014; Denais et al., 2016), with a migration chamber height of 5 µm and a pore width of either 1, 2 or 15 µm, resulting in 5, 10 and 75 µm^2^ cross-sectional areas. In brief, the migration devices were coated with rat tail type-I collagen (50 µg/mL; BD Biosciences) in acetic acid (0.02 N) or human plasma fibronectin (5 µg/mL; Millipore) in PBS overnight at 4°C and rinsed with imaging medium to remove the coating solution. After loading the cells, devices were incubated for at least 3 hours to allow cell attachment before imaging. Microfluidic migration devices were imaged at 37°C on an inverted Zeiss Observer Z1 microscope equipped with a CCD camera (Photometrics CoolSNAP KINO) using a 20×/NA 0.8 air objective. The image acquisition was automated through ZEN (Zeiss) software with imaging intervals of 5 min collecting images for DIC and GFP. Time-series images were stabilized using a custom written MATLAB (Mathworks) script that used features of the PDMS device as fiducials to compensate for the inaccuracy of the linear encoded microscope stage. Individual cells were tracked during the process of pore transmigration by using a custom written Matlab script (Elacqua et al., 2018) with manual verification.

### Confocal fluorescence and reflection microscopy and image analysis

For fixed-cell microscopy, cells were cultured on glass for 24 h, or in polymerized collagen for 0-20 h before fixation with 2% or 4% buffered PFA or methanol, permeabilized with 0.1% Triton100, and stained with the indicated primary and secondary antibodies and dyes. Samples were monitored by confocal laser scanning microscopy (Zeiss LSM 510, 40×/NA 1.3 oil objective and 40x/NA 0.8 water objective; Olympus, 20x/NA 0.5 water objective and 40x/NA 0.8 water objective) with simultaneous 3-5 channel scanning for fluorescence, reflection and transmission signal, and as z-stacks of 1-2 μm distance. For life-cell microscopy, cells were monitored by a BD Pathway 855 high content microscope with spinning disk (20×/NA 0.75 air objective), or a Leica SP5 laser scanning microscope using a temperature- and CO_2_ -controlled stage (37°C, 5%).

Image analysis was performed by Fiji ImageJ (software version 1.40v; National Institutes of Health): XY image stacks were either displayed as single scan, reconstructed as maximal projections, or reconstructed as XZ images using the reslice function, as indicated in legends. In still images, elongation index (length/width) and the smallest nuclear diameter of cells were calculated from images after indicated hours of migration within collagen. Round non-moving, apoptotic or mitotic cells, as identified morphologically by actin signal, or from the movie sequence and transmission signals, were excluded from analysis. Image processing was performed by cropping, rotation, and manual adjustment of contrast and brightness and displayed in virtual colors.

Dynamic changes in nuclear shapes over time were analyzed as in (Krause et al., 2019). In brief, image stacks were thresholded, nuclear outlines generated, and nuclear fluctuation [as adapted from (Lee et al., 2014)] calculated. Fluctuation was calculated by overlaying the two consecutive images with their centroids and performing a 1°-stepwise full-circle (360°) rotation for the maximum overlap between these two shapes. The resulting fluctuation number was computed through

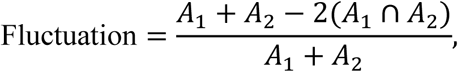

Where *A*_1_, *A*_2_ are the areas of the two consecutive nuclear shapes (monitored 4 min after eachother) and *A*_1_ ∩ *A*_2_ are the maximum intersections found through the rotation.

### Statistics

Significance was determined by the two-tailed unpaired Students t-test (western blots, transmigration rates, NLS leakage), or a two-tailed unpaired Mann-Whitney test for non-gaussian distributed cell samples (image analyses, AFS measurements, migration rates), respectively, using the GraphPad Prism 5 program. In order to compare data from individual experiments (n’s), one median from all the respective control cell measurements was calculated, and a correction factor applied on the data of the individual experiment (see detailed explanation in legend of Fig. S2).

## Online supplemental material

**Fig. S1** shows lamin levels in populations of HT1080 cells after modulation of lamin isoforms.

**Fig. S2** describes the principles and measurements from lamin-modulated HT1080 cells derived by atomic force microscopy.

**Fig. S3** shows data generated from additional cell lines MV3 melanoma, MDA-MB-231 and MCF10A breast cancer cells after down- and upregulation of lamin isoforms.

**Fig. S4** depicts collagen pore size ranges from different collagen sources, and HT1080 cell migration data in collagen after lamin modulation.

**Fig. S5** shows data on HT1080 and MDA-MB cell migration after lamin modulation in transwell chamber and microchannel assays, as well as NE rupture incidence.

**Video 1** shows compression and decompression of the nucleus from a GFP-lamin A expressing HT1080 cell by a bead coupled cantilever.

**Video 2** shows aspiration of HT1080 cells and their colored nucleus into a micropipette by the application of high pressure.

**Video 3** shows a confocal microscopy-derived time sequence of a single HT1080 cell migrating in confining collagen and adapting its nucleus labeled with H2B-mCherry and GFP-lamin A.

**Video 4** shows the effect of lamin isoform downregulation on HT1080 cell migration rates in confining collagen.

**Video 5** shows the effect of lamin isoform overexpresssion on HT1080 cell migration rates and cell morphology in confining collagen.

**Video 6** shows nuclear pore passage time of HT1080 cells after lamin isoform downregulation in a snythetic microdevice.

**Video 7** shows the effect of lamin A overexpresssion on MDA-MB-231 cell migration rates and cell morphology in confining and optimal collagen pore conditions.

**Video 8** shows the nuclear rotation method for analysis of nuclear fluctuation.

**Video 9** shows segmented nuclear shapes and their dynamic changes in HT1080 cells migrating in confining collagen after downregulation of lamin isoforms.

**Figure legends, Supplementary figure legends, Video legends:** see Figure part

---

**Figure S1.**
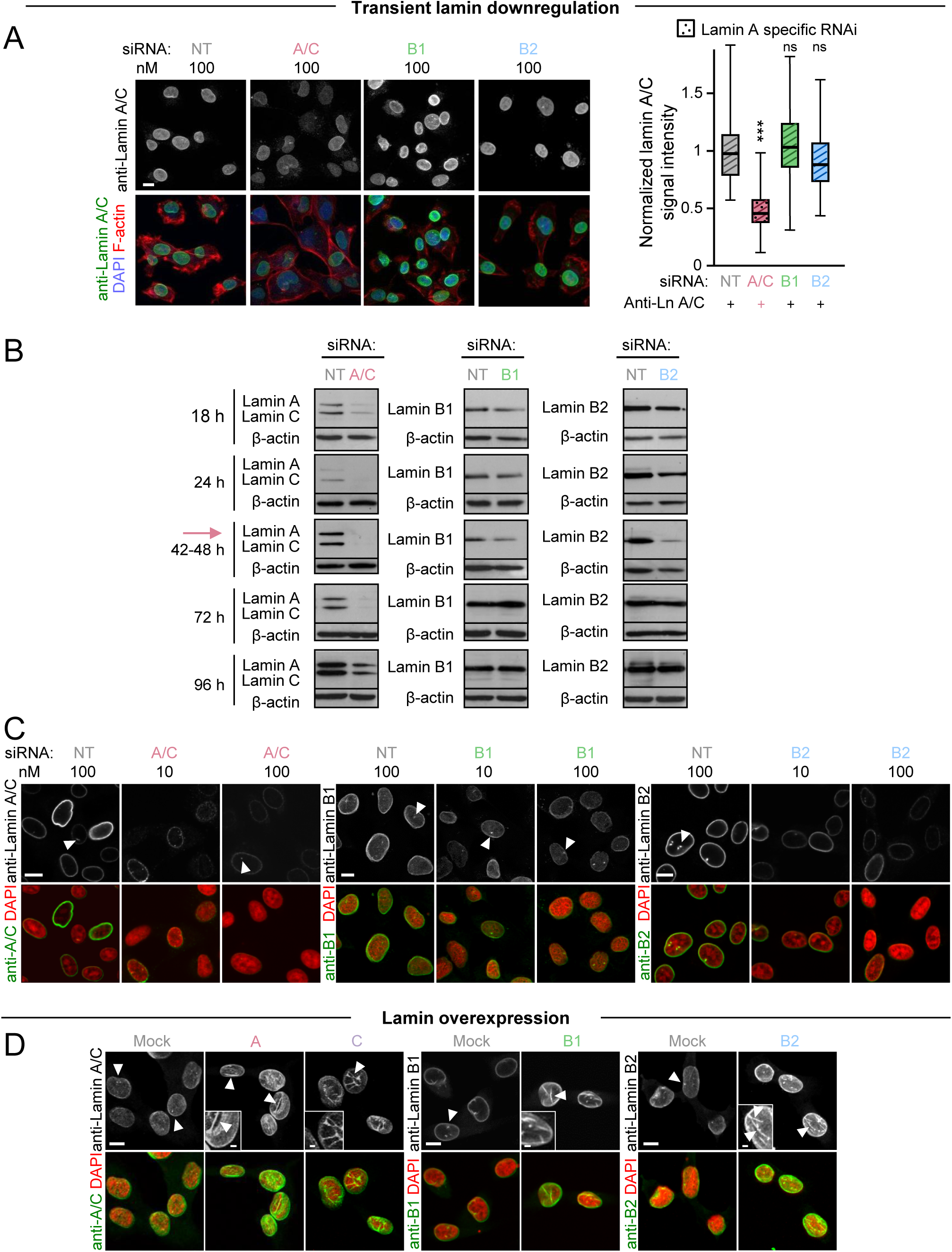
HT1080 cell lamin levels after expression modulation. Nuclear morphology and lamin localization and intensity after transient lamin downregulation or stable overexpression **(A,C,D)**. 2D cell cultures on glass were fixed by PFA (A,D) or methanol (C), stained and imaged by confocal microscopy. Images are displayed as single central Z-scans. **(A)** Lamin A/C intensity after transient downregulation by lamin-isoform-specific siRNA of each 100 nM; striped bars, as in all following data sets). Confocal imaging (left) and graph (right) for mean lamin intensity per nucleus (n = 1; 40-66 cells per condition), as determined from images as shown left, but quantified from projections as shown in Fig. 1A. Black lines, boxes and whiskers; medians, 25^th^/75^th^, and 5^th^/95^th^ percentile. For statistical analyses, non-paired Mann-Whitney test was used. **(B)** Changes of lamin levels over indicated time periods after lamin downregulation; determined by western blot. Sample lysates were generated after indicated time points post transfection with 100 nM siRNA. Lamin expression was found to be most reduced after 42-48 h (red arrow), and was 12-13% for lamin A/C, 52% for lamin B1 and 20% for lamin B2, as compared to 100% lamin expression in non-targeting siRNA treated cells and normalized to β-actin protein content. **(C)** Representative images of lamin downregulation 48 h after transfection with 10 or 100 nM siRNAs targeting indicated lamin isoforms. White arrowheads indicate intranuclear speckles. **(D)** Stable overexpression of indicated lamin isoforms, 24 h after cell seeding. White arrowheads indicate intranuclear speckles/ tubes or membrane foldings, respectively (also see insets). All image bars: 10 μm, except insets (2μm).

**Figure S2.**
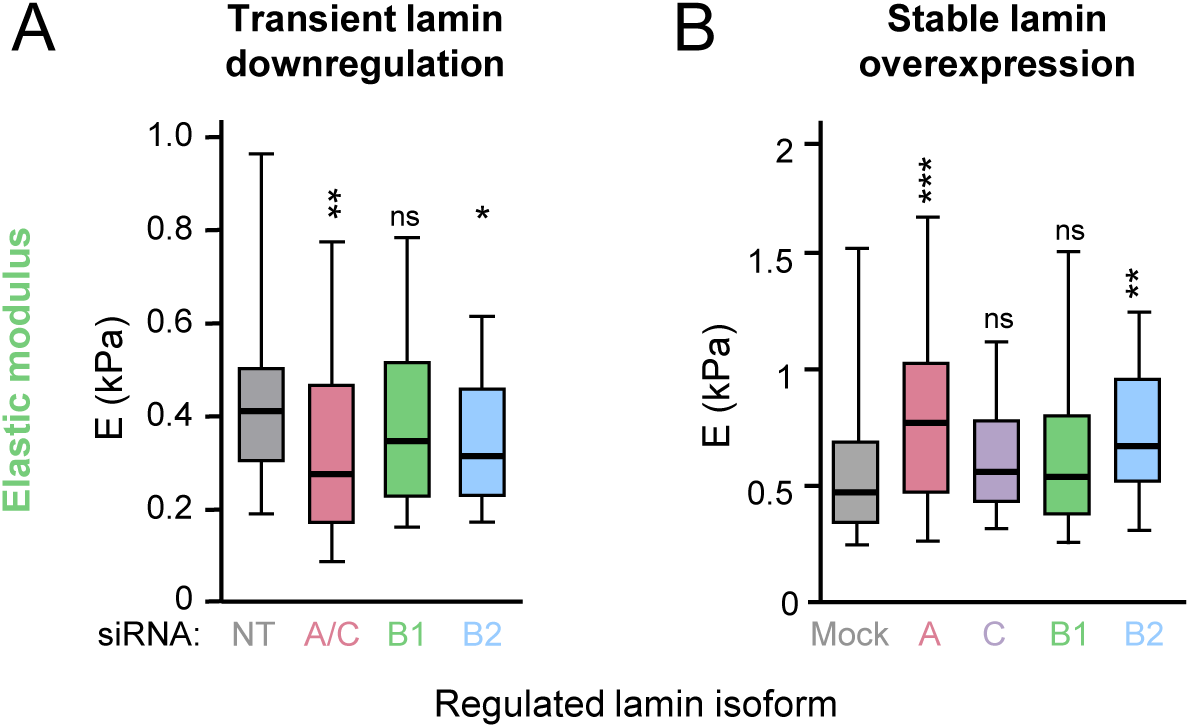
AFS-derived elastic modulus of HT1080 cells after modulation of lamin expression. HT1080 cells were either treated by specific indicated siRNA’s (10 nM; **A**) or stably overexpressed with lamin-specific or control vectors **(B)**. Elastic modulus E of cells after **(A)** lamin A/C, B1 and B2 siRNA treatment calculated from 0.5 nN force curves; n = 2-4; 28-66 cells per condition, and **(B)** stable lamin overexpression calculated from 2 nN force curves; n = 2-5; 23-48 cells per condition. To better compare experiments carried out on different days, medians from the individual non-targeted control cell populations were calculated (i.e. 0.50; 0.58; 0.62; 0.50). Then one median (i.e. 0.53) was formed based on the data of all control cells. Subsequently, a correction factor for each experiment was calculated (i.e.1.06; 0.91; 0.86; 1.06) and applied for each measurement. Black horizontal lines, boxes and whiskers show the medians, 25^th^/75^th^, and 5^th^/95^th^ percentile. For all experiments, ***, P <0.001; **, P <0.01; *, P <0.05; ns, and non-significant (non-paired Mann-Whitney test).

**Figure S3.**
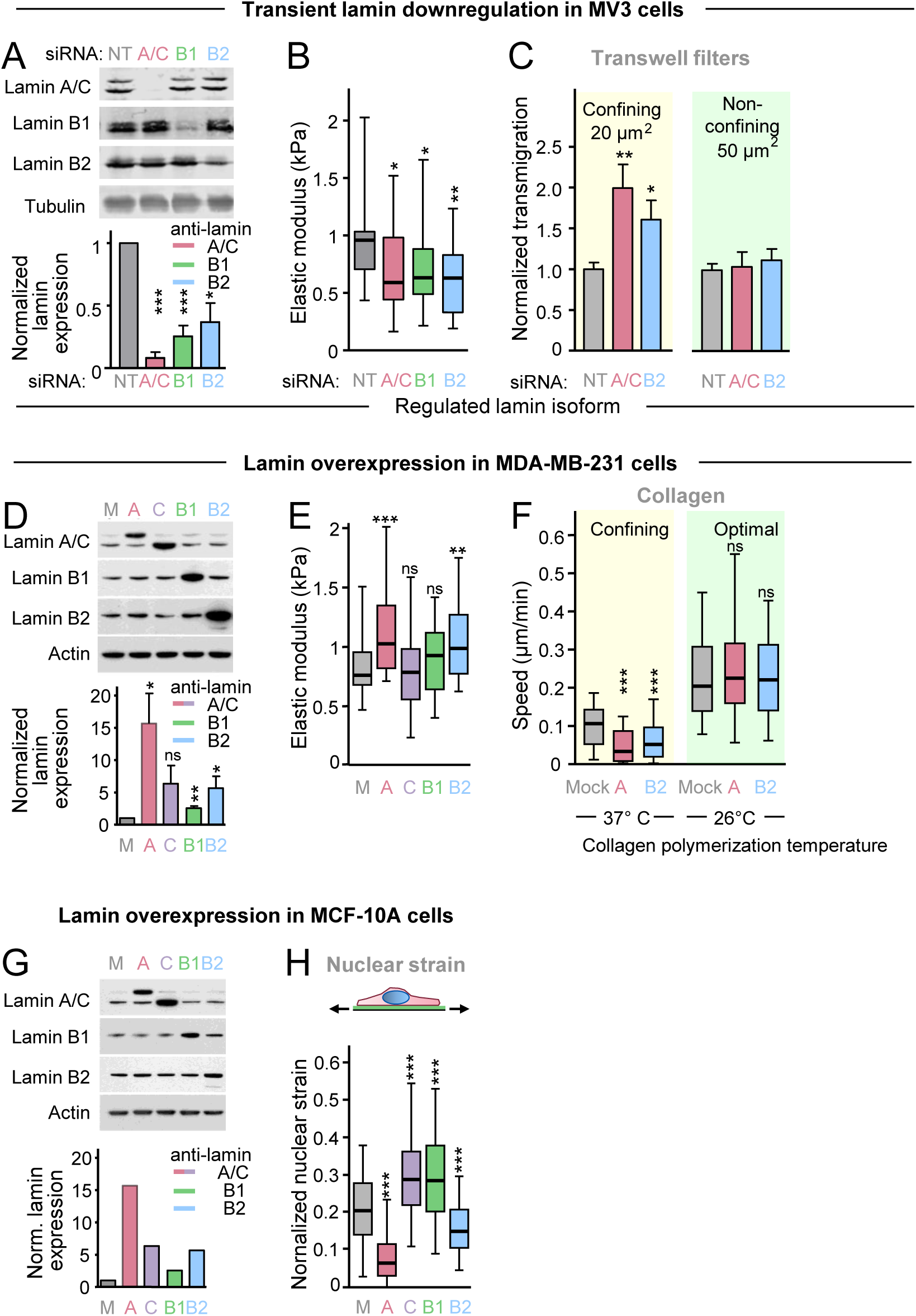
Measurements of lamin-mediated nuclear elasticity and migration rates in different cell types. MV3 melanoma cells were transfected with 10 nM indicated lamin-specific siRNAs before the experiment **(A-C)**, and MDA-MB-231 **(D-F)** and MCF10A cells **(G-H)** were stably transduced with vectors coding for indicated lamin isoforms. **(A,D,G)** Top, representative western blots for lamin A, C, B1 and B2 levels and tubulin or actin loading control from whole cell lysates. Bottom, expression levels of specifically regulated lamin isoforms after calculation of signal intensity by densitometry and normalization to both loading control and control siRNA transfection. M, Mock vector transduced control cells. Means and SEM are shown; n = 3 **(A,D)**; n=1 **(G). (B)** Elastic modulus of cells after indicated siRNA treatment. Cells were probed with 1.5 nN applied force and values were calculated into E in kPa. N = 2; 22-30 cells per condition. **(C)** Normalized transmigration rates of cells that have successfully passed to the lower side of a 5 µm or 8 µm porous membrane, respectively. N = 1-3; bars represent mean and whiskers are SEM. **(E)** Elasticity measurements, calculated from 2 nN force; n = 2-3; 30-35 cells per condition. **(F)** Averaged single cell migration rates over 24 h in collagen lattices of bovine origin polymerized at indicated temperatures (see Fig. 6D) and in the presence of GM6001; n = 1-3, 28-107 cells per condition. **(H)** Top, cartoon depicting the principle of nuclear strain measurements. Bottom, nuclear strain measurements from MCF10A cells adherent to… N = ≥3; 124-157 cells per condition. **(A,C,D,G)** Bars and whiskers, mean and SEM; **(B,E,F,H)** horizontal lines, boxes and whiskers showing the medians, 25^th^/75^th^, and 5^th^/95^th^ percentile. For all experiments, ***, P <0.001; **, P <0.01; *, P <0.05; ns, non-significant; **(A,C,D)** two-tailed unpaired Students t-test); **(B,E,F,H)** non-paired Mann-Whitney test.

**Figure S4.**
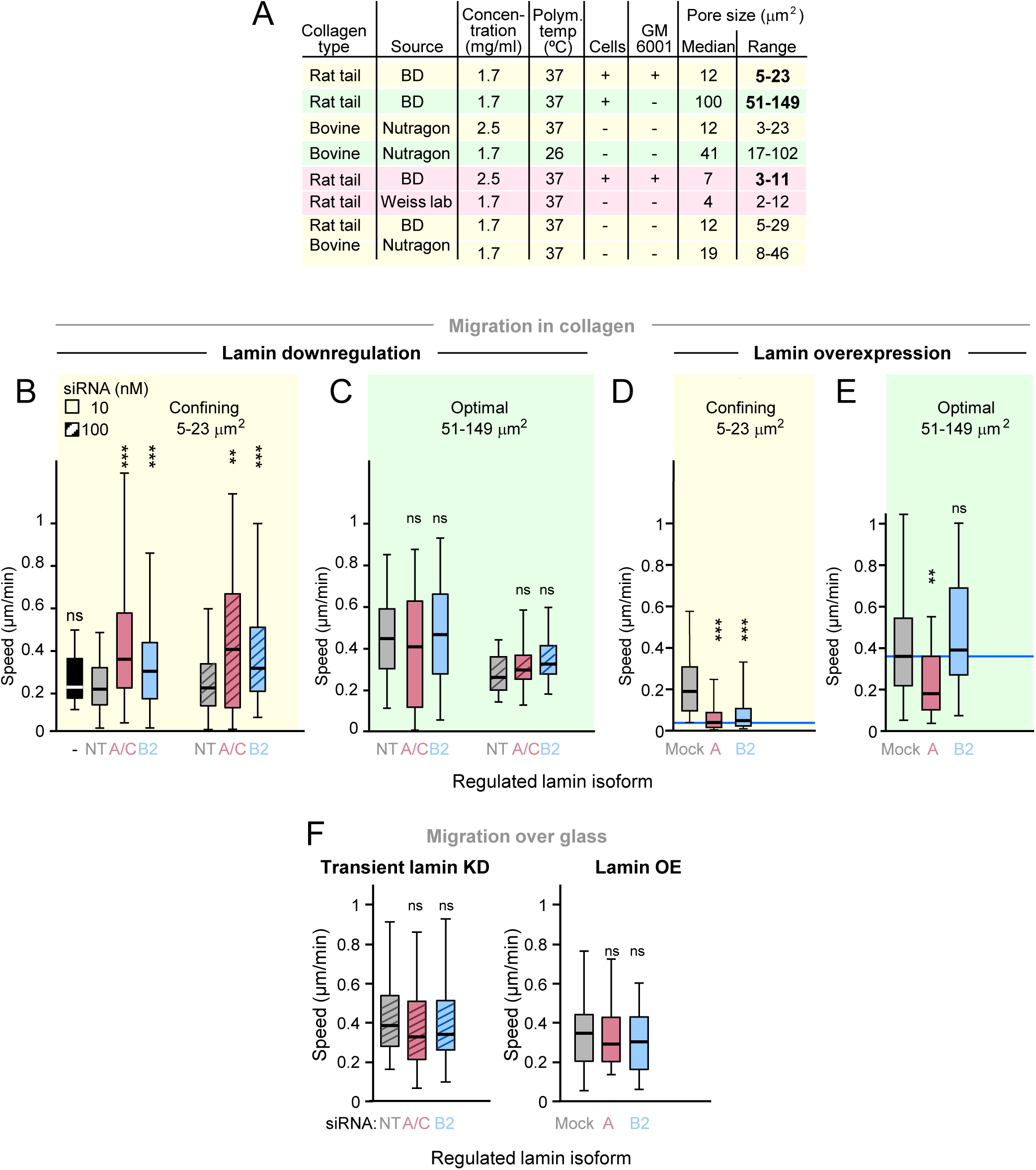
Impact of lamin modulation on HT1080 migration rates in collagen of varying porosity or on glass. **(A)** Comparison of pore areas quantified from different indicated collagen sources and densities (see Denais et al., 2016; Wolf et al., 2013); 21 data points per condition. Colored areas indicate confining (yellow), non-confining (green) and (near-) limiting (red) pore conditions induced by different collagen concentrations, sources, and by the presence or absence of proteolysis during cell migration. **(B-E)** Cell migration in 3D collagen of indicated porosity monitored by time-lapse microscopy over 20-24 h. **(B,C)** Averaged single cell migration rates after transient downregulation by indicated concentrations of specific siRNA, analyzed by single-cell tracking. Cells migrated in rat tail collagen (1.7 mg/ml) in the presence **(B)** or absence **(C)** of GM6001. **(D,E)** Averaged single cell migration rates after stable overexpression of control or indicated lamin vectors. Cells migrated in rat tail collagen (1.7 mg/ml) in the presence **(D)** or absence **(E)** of GM6001. **(F)** Cell migration over the surface of a cell culture dish. Cells were monitored by time-lapse microscopy and analyzed by cell tracking. Averaged single cell migration rates after treatment with 100 nM indicated specific siRNA’s; or after stable expression by empty vector or indicated lamin isoforms. **(B-F)** Horizontal lines, boxes and whiskers show the medians, 25^th^/75^th^, and 5^th^/95^th^ percentile. **(D,E)** Horizontal blue lines mark 10% migration rates (0.036 μm/min) in confinement as compared to migration of Mock cells in optimized conditions (0.36 μm/min). **(B)** N = 2-5, 52-166 cells; **(C)** n = 2-3, 16-31 cells; **(D)** n = 3; 87-93 cells; **(E)** n = 1-2, 30-62 cells; **(F)** n = 2, 44-61 cells (left graph); n = 2; 66-71 cells per condition (right graph). ***, P <0.001; **, P <0.01; *, P <0.05; ns, non-significant; by non-paired Mann-Whitney test.

**Figure S5.**
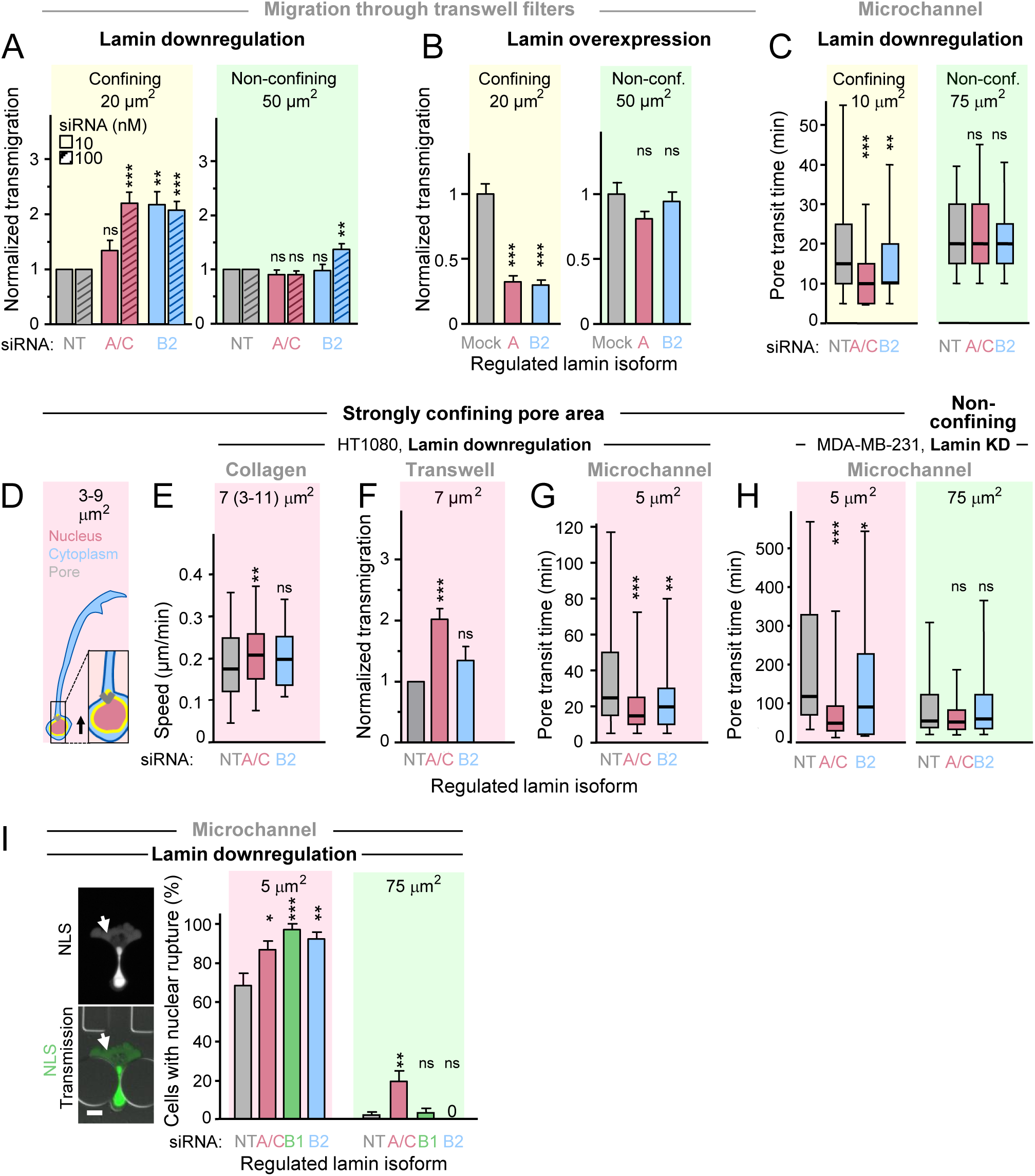
Cell migration and nuclear envelope rupture in synthetic substrates at non-confining, confining and limiting pore conditions. HT1080 **(all, except H)** and MDA-MB-231 **(H)** cells were modified for lamin expression as indicated and allowed to migrate through transwell membranes **(A,B,F)**, microchannel devices **(C,G,H,I)**, or collagen **(E). (A,B,F)** Depiction of transmigration rates of cells that have successfully transmigrated to the lower side of the transwell membrane through pores at indicated cross-sections (see Fig. 4A). Values were normalized towards means of transmigrated control cell numbers at indicated conditions (prior to normalization, after applying 10.000 cells to the upper compartment, approximately 2500, 1200 and 600 cells transmigrated through the 8, 5 and 3 µm transmembrane filters, respectively). N = 2-7. **(C,G,H)** Cell migration through through a 3D PDMS microchannel device (see Fig. 4C). Depicted is the time for colored cell nuclei to pass indicated pore cross sections. **(C,G)** N = 3; 53-61 HT1080 cells per condition; **(H)** n = 3; 23-50 MDA-MB-231 cells per condition. **(I)** Left, images of cells with NLS leakage into the cytoplasm (arrow) indicative of nuclear envelope rupture in the moment of transmigration through a 5 µm^2^ constriction. Bar, 10 μm. Right, incidence of NE rupture after lamin A/C, B1 or B2-specific RNAi, or nontarget (NT) control. N = 3; 36-60 cells per condition. **(D)** Depiction of cell and nuclear morphology at limiting pore size (red background coloring; Wolf et al., 2013). Nuclei are round or contain occasional thin protrusions with a deformation limit of ≤3 μm diameter or around 7 μm^2^ cross section that cannot be exceeded further (Wolf et al., 2013). Likewise, migration-abrogated cells form long cytoplasmic protrusions. Nearly abrogated cell locomotion rate is depicted by short arrow length (compare with Figure 3A). **(E)** Mean single cell migration rates in rat tail collagen (2.5 mg/ml) in the presence of GM6001; n = 1-4, 8-141 cells. **(A,B,F,I)** Bar diagrams, mean + SEM. **(C,E,G,H)** Box plot diagrams; horizontal lines, boxes and whiskers show the medians, 25^th^/75^th^, and 5^th^/95^th^ percentile. ***, P <0.001; **, P <0.01; *, P <0.05; ns, non-significant. **(A,B,F,I)** Two-tailed unpaired Students t-test; **(C,E,G,H)** non-paired Mann-Whitney test. KD, knockdown.

